# Analysis of *Epichloë festucae* small secreted proteins in the interaction with *Lolium perenne*

**DOI:** 10.1101/490425

**Authors:** Berit Hassing, David Winter, Yvonne Becker, Carl H. Mesarich, Carla J. Eaton, Barry Scott

## Abstract

*Epichloë festucae* is an endophyte of the agriculturally important perennial ryegrass. This species systemically colonises the aerial tissues of this host where its growth is tightly regulated thereby maintaining a mutualistic symbiotic interaction. Recent studies have suggested that small secreted proteins, termed effectors, play a vital role in the suppression of host defence responses. To date only a few effectors with important roles in mutualistic interactions have been described. Here we make use of the fully assembled *E. festucae* genome and EffectorP to generate a suite of 141 effector candidates. These were analysed with respect to their genome location and expression profiles *in planta* and in several symbiosis-defective mutants. We found an association between effector candidates and a class of transposable elements known as MITEs, but no correlation with other dynamic features of the *E. festucae* genome, such as transposable element-rich regions. Three effector candidates and a small GPI-anchored protein were chosen for functional analysis based on their high expression *in planta* compared to in culture and their differential regulation in symbiosis defective *E. festucae* mutants. All three candidate effector proteins were shown to possess a functional signal peptide and two could be detected in the extracellular medium by western blotting. Localization of the effector candidates *in planta* suggests that they are not translocated into the plant cell, but rather, are localized in the apoplastic space or are attached to the cell wall. Deletion and overexpression of the effector candidates, as well as the putative GPI-anchored protein, did not affect the plant growth phenotype or restrict growth of *E. festucae* mutants *in planta*. These results indicate that these proteins are either not required for the interaction at the observed life stages or that there is redundancy between effectors expressed by *E. festucae*.

## Introduction

Plant-pathogenic fungi deploy a number of virulence factors, termed effector proteins, to promote colonization of their hosts. These effector proteins, which can be targeted to the host apoplast (extracellular effectors) or to various compartments of the plant cell (intracellular effectors), typically promote colonization by altering host physiology or by modulating host immune responses [1]. Well-characterized examples of extracellular fungal effector proteins include Avr2, Avr4 and Ecp6 from the tomato leaf mould pathogen *Cladosporium fulvum* [2-4], as well as Pit2 and Rsp3 from the corn smut pathogen *Ustilago maydis* [5, 6]. More specifically, Avr4 binds to chitin molecules present in the cell wall of invading hyphae to prevent hydrolysis by host chitinases [2, 7], while Ecp6 sequesters chitin oligosaccharides released from the cell wall of invading hyphae to prevent detection by host chitin immune receptors [4, 8]. Rsp3 binds to the cell wall of invading hyphae to protect them against two antimicrobial mannose-binding host proteins [6], while both Pit2 and Avr2 inhibit host cysteine proteases to prevent degradation of fungal proteins [3, 9-11]. A well-characterized example of an intracellular fungal effector protein is Cmu1 from *U. maydis*, which functions as a chorismate mutase to redirect the metabolism of chorismate away from the production of the defense signaling hormone salicylic acid [12].

Like plant-pathogenic fungi, plant-beneficial fungi must also deploy effector proteins to promote host colonization [13, 14]. However, to date only a small number of effectors from this group of fungi have been identified and functionally characterized. Furthermore, of those that have been functionally characterized all are from mycorrhizal fungi. One such example is MiSSP7 from *Laccaria bicolor*, an ectomycorrhizal fungus of poplar roots. In plants (including poplar), JAZ proteins repress the expression of jasmonic acid (JA)-related defense genes, with the degradation of these proteins leading to the activation of JA-related defense genes [15]. During infection of poplar, MiSSP7 is transported into the plant nucleus where it interacts with and stabilizes the JAZ protein PtJAZ6 to prevent its degradation, and thus prevent the expression of JA defence-related genes [16, 17]. A second example is SP7, which is produced by the arbuscular mycorrhizal fungus, *Glomus intraradices*. SP7, which like PtJAZ6 is also transported into the plant nucleus, influences defense gene signaling by interacting with the pathogenicity-related transcription factor ERF12 [18]. A third example is Fgb1 has been characterized from the endophytic root-colonizing fungus, *Piriformospora indica*. Fgb1 is a β-glucan-binding lectin that modulates the fungal cell composition and prevents β-glucan-triggered immunity in plants, presumably through a similar mechanism to that shown for as Ecp6 and other LysM domain-containing effectors from plant-pathogenic fungi [19].

There is little amino acid sequence conservation between effector proteins of plant-associated fungi, and most lack obvious functional domains. Together, these features render their identification and functional characterization difficult. Some features, however, are common to many effector proteins. For example, most have an amino (N)-terminal signal peptide for secretion, are small (<300 amino acid residues in length) and rich in an even number of cysteines, and are highly expressed during host colonization [1, 20]. Furthermore, many of these effector proteins are encoded by genes located in dynamic regions of the fungal genome (e.g. those regions rich in transposable elements; TEs), where increased mutation rates can result in the increased diversification of effector genes; a beneficial trait in the ‘arms race’ with the host [1, 21, 22]. One group of TEs are miniature inverted-repeat transposable elements (MITES)[23], which are small non-autonomous DNA transposable elements that have been proposed to have a role in regulating the expression of plant induced genes in phytopathogenic and mutualistic fungi [24-27].

Here we present an analysis of effector candidates from *Epichloë festucae*, an endophytic fungus of the cool-season grass, *Lolium perenne*. The interaction between *Epichloë spp.* and cool-season grasses is of agricultural importance due to the array of secondary metabolites produced by the endophyte that protect the host from various biotic stresses, such as insect and mammalian herbivory [28]. Unlike mycorrhizal fungi, which only infect the roots of their host plants, *E. festucae* and other *Epichloe* spp. exclusively colonize plant aerial tissues and are only found in association with their host. These endophytes can be vertically transmitted through host seeds [29, 30] or horizontally transmitted by ascospore transfer. The latter occurs following sexual development on host inflorescences, and results in the formation stromata that prevent the maturation of these inflorescences to cause ‘choke’ disease [31, 32]. While some *Epichloë* spp. have a high degree of host specificity, others, such as *E. typhina,* can infect a variety of grass species [33]. Within the host tissue, the fungal growth is highly restricted and confined to the intercellular spaces. Deletion of key signaling genes results in the loss of this restricted growth and, in doing so, the ‘breakdown’ of the symbiosis [34-40]. Here, we make use of the fully assembled *E. festucae* genome [26] and EffectorP [41] to identify a suite of candidate effectors in this mutualistic endophyte, and analyse their genomic location to identify potential patterns of distribution. In addition, we use multiple transcriptome datasets to generate an expression profile for these candidate effectors, with the aim of identifying effectors with a potential role in the *E. festucae*-*L. perenne* interaction. Based on these analyses, three candidate effectors were selected for functional characterization. As GPI-anchored proteins play an important role in plant– fungus interactions [42, 43] we also selected a GPI anchored protein with a similar expression file to the abovementioned candidate effectors for functional characterization.

## Materials and Methods

### Bioinformatics

A detailed record of the bioinformatic and statistical analyses performed in this study is provided in S1 File. Here we briefly describe each step of these analyses.

#### Classification of proteins and amino acid composition

The program SignalP v4.1 [44] was used to identify those *E. festucae* proteins of ≤200 amino acid residues in length that contain an N-terminal secretion signal, while EffectorP v 1.0 [41] was used to predict which of these proteins is an effector. The latter uses a machine learning approach based on sequence properties common to known effectors. Taking these results together, each protein was then classified into one of three non-overlapping classes: (i) candidate effectors (proteins with ≤200 amino acid residues, a signal peptide and an EffectorP probability of >0.5), (ii) secreted proteins (proteins with a signal peptide that do not meet the other abovementioned criteria for effectors), and (iii) non-secreted proteins (all other proteins). The hypothesis that secreted proteins are more cysteine-rich than average was tested using a logistic regression. Specifically, a model with the proportion of cysteine residues as the response variable and membership of the three protein-classes described above as a predictor was fitted using R 3.0.1 [45].

### Genomic landscape of candidate effectors

Bedtools v 2.26.0 [46] and the recently published *E. festucae* Fl1 genome [26], were used to calculate the distance between each protein-coding gene and its nearest AT-rich isochore, minature inverted TE (MITE) and telomere. Logistic regression was used to test whether secreted proteins and candidate effectors are more likely to appear near to a MITE (within 2 kb), an AT-rich isochore (within 5 kb) or a sub-telomeric region (within 50 kb) than other proteins. A Monte Carlo approach was used to test whether genes encoding candidate effectors fall into clusters within the Fl1 genome. Specifically, Bedtools was used to calculate the minimum distance between each effector-encoding gene and another gene of the same type (i.e. the minimum distance to another candidate effector gene). The mean distance between candidate effector genes was compared to a null distribution of this statistic. We simulated this null distribution by calculating the same statistic for 1000 random gene-sets, each the same size as the set of candidate effectors.

### Gene expression analyses

The hypothesis that genes encoding candidate effectors are differentially expressed *in planta* compared to axenic culture was tested by logistic regression, using previously published RNAseq data as input [36, 47]. A model in which the probability that genes demonstrate significant (corrected p-value < 0.01) and substantial (> 2 log_2_ fold) differences in expression between culture and *planta* conditions compared between candidate effectors, secreted proteins and all other proteins was fitted using R. Similarly, the hypothesis that repression of candidate effector genes is disrupted in Δ*hepA*, a mutant of the heterochromatin protein 1 (HP1) gene [48, 49], was tested using logistic regression. Finally, the *in planta* expression of candidate effector genes was compared across four separate symbiosis-deficient mutants, including Δ*noxA*, Δ*proA* and Δ*sakA* [36, 47] and Δ*hepA* (Chujo & Scott, personal communication. The Δ*hepA* transcriptome data used here is available from the Sequence Read Archive (SRA) under Bioproject PRJNA447872. A list of the individual biosample numbers is provided in S1 Table.

### Strains and growth conditions

*Escherichia coli* strains were grown overnight in Lysogeny Broth (LB) or on LB agar supplemented with 100 µg/ml ampicillin or 50 µg/ml kanamycin at 37°C, as described by [50]. *Agrobacterium tumefaciens* strains were grown for one day in LB broth or for two days on LB agar supplemented with 50 µg/ml kanamycin at 30°C, as previously described [51]. *Saccharomyces cerevisiae* strains were grown at 30°C on YPDA (2% peptone, 1% yeast extract 0.01% adenine hemisulfate, 2% glucose, 2% agar; adjusted to pH 5.7). For *S. cerevisiae* complementation assays cells were plated on CMD tryptophan dropout medium (0.67% yeast nitrogen base, 0.064% W/L dropout supplement, 0.00033% leucine, 2% sucrose, 0.1% glucose, 2% agar, pH 5.7). To test for secretion of the SUC2 secretion signal fusion protein, fresh colonies of *S. cerevisiae* were streaked on CMDRAA medium (0.67% yeast nitrogen base, 0.064% W/L dropout mixture, 0.00033% leucine, 0.00033% tryptophan, 2% raffinose, 2 µg/ml antimycin A, 2% agar, pH 5.8) and compared with growth on YPDA control plates [52, 53]. *E. festucae* strains were grown on 2.4% Potato-Dextrose Agar (PDA) supplemented with 150 µg/ml hygromycin B or 250 µg/ml geneticin or in 2.4% Potato-Dextrose Broth (PDB) for up to seven days [54, 55]. For microscopy analyses, strains were grown on 3% water agar overlaid with 1.5% water agar [40]. All strains used in this study can be found in S2 Table.

### Plant growth and endophyte inoculation

*E. festucae* strains were inoculated into endophyte-free seedlings of perennial ryegrass (*L. perenne* cv Samson) using the method previously described [56]. Inoculated plants were grown in root trainers at 22°C in a controlled growth room with a photoperiod of 16 h light (approximately 100 μE/m^2^/s) and 8 h dark. *N. benthamiana* and *N. tabacum* were grown at approx. 28°C in a controlled growth room with a photoperiod of 16 h light (approximately 100 μE/m^2^/s) and 8 h dark for 6–7 weeks. For agroinfiltration experiments (see below), plants were transferred to a greenhouse and grown at 25°C with natural lighting.

### DNA isolation, PCR and sequencing

High quality genomic DNA was extracted from freeze-dried mycelium of *E. festucae* as previously described [57]. Plasmid DNA was extracted from *E. coli* using the High Pure plasmid isolation kit (Roche). DNA resolved by agarose gel electrophoresis was purified using the Wizard SV Gel and PCR Clean-Up System (Promega) or the Zymoclean Gel DNA Recovery kit (Zymo Research). PCR amplification of short DNA products was conducted with the OneTaq^®^ (NEB) polymerase according to the manufacturer’s instructions. Each reaction was set up as follows: 1× Standard OneTaq Reaction Buffer, 200 µM dNTPs, 0.2 µM for each forward and reverse primer, 0.025 units/µl OneTaq Polymerase and 20-300 ng of template DNA. If crude DNA was used as a template, 3% DMSO was added. PCRs were performed with the following Thermocycler (Eppendorf) settings: 1 min at 94°C for initial denaturation, then 30-35 cycles of 30 s at 94°C (denaturation), 30 s at 45-65°C (annealing), 1 min/kb at 68°C (extension) followed by 3-5 min at 68°C (final extension). Amplification of DNA fragments for Gibson Assembly [58] was performed with Phusion High Fidelity polymerase (ThermoFisher Scientific). Each reaction was set up as follows: 1× Phusion HF buffer, 200 µM dNTPs, 0.5 µM of each forward and reverse primer, 0.02 U/µl Phusion polymerase and 20-300 ng of template DNA. PCRs were performed with the following Thermocycler settings: 1 min at 98°C for initial denaturation, then 30-35 cycles of 30 s at 98°C (denaturation), 30 s at 59-68°C (annealing), 30 s/kb at 72°C (extension), followed by 3-5 min at 72°C (final extension). Where long primers (>30 bp) were used, a 2-step PCR was performed with the same reaction set up, but with different cycler settings: 1 min at 98°C for initial denaturation, then 5 cycles of 30 s at 98°C (denaturation), 30 s at 55°C (annealing), 30 s/kb at 72°C (extension), followed by 25 cycles of 30 s at 98°C (denaturing) and 30 s/kb at 72°C (extension). Subsequently the reaction was incubated for 3-5 min at 72°C (final extension). PCR products were purified using the Wizard SV Gel and PCR Clean-Up System (Promega). Primers were sourced from Integrated DNA Technologies. Primer sequences can be found in the Supplemental Table S2. Sequencing reactions were performed using the Big-DyeTM Terminator Version 3.1 Ready Reaction Cycle Sequencing Kit (Applied BioSystems, Carlsbad, California, USA), and separated using an ABI3730 genetic analyser (Applied Bio Systems). Sequence data was assembled and analysed using MacVector sequence assembly software, v14.5.2.

### Preparation of constructs

The *gpiB* replacement construct, pBH1, was prepared as follows. Firstly, the 1,639 bp 5′ and 1,389 bp 3′ flanks of *gpiB* were amplified from wild type (WT) *E. festucae* strain Fl1 genomic DNA by PCR using BH1/BH2 and BH3/BH4 primer pairs, respectively. The 1,394 bp hygromycin resistance cassette (P*trpC*-*hph*) sequence was then amplified from pSF15.15 DNA using the hph_F/hph_R primer pair. The 5,483 bp backbone vector, pRS426, was amplified by PCR using the pRS426_F/pRS246_R primer pair. All PCRs were performed with Phusion High-Fidelity DNA polymerase (NEB). The fragments were then assembled via Gibson Assembly [58] and the correct assembly was verified using a diagnostic restriction enzyme digest and sequencing. The replacement constructs for *sspM* (pBH2), *sspN* (pBH3) and *sspO* (pBH4) were generated in the same way. The 1,377 bp 5’ and 1,333 bp 3’ flanks of *sspM* were amplified with the BH11/BH12 and BH9/BH10 primer pairs, respectively; the 1,431 bp 5’ and 1,433 bp 3’ flanks of *sspN* were amplified with the BH15/BH16 and BH13/BH14 primer pairs, respectively; and the 1,443 bp 5’ and 1,318 bp 3’ flanks of *sspO* were amplified with the BH5/BH6 and BH7/BH8 primer pairs, respectively.

The *S. cerevisiae* complementation vector pSUC2T7M13ORI [52] was fully sequenced using primers BH25 to BH42 and BH61 to BH63. The nucleotide sequences encoding the secretion signals from SspN (54 bp), SspO (54 bp), SspM (54 bp) and GpiB (57 bp) were then amplified from *E. festucae* strain Fl1 genomic DNA with the BH21/BH22, BH70/BH71, BH68/BH69 and BH64/BH65 primer pairs, respectively, using OneTaq polymerase. In doing so, these primers added a 5’ *Eco*RI and 3’ *Xho*I restriction site to each amplified sequence. Due to difficulties in working with small fragments different cloning strategies were effective for the different fragments. The *sspN* fragment was cloned into the TOPO-TA vector according to the manufacturer’s instructions. Following this cloning step, pSUC2TM13ORI and the TOPO-TA vector containing the *sspN* secretion signal-encoding nucleotide sequence were digested with *Eco*RI and *Xho*I restriction enzymes and purified. The pSUC2TM13ORI vector backbone and secretion signal-encoding nucleotide sequence were then ligated using T4 ligase to generate pBH7. The *gpiB, sspM* and *sspO* secretion signal-encoding nucleotide sequences were directly digested with *Eco*RI and *Xho*I and purified, omitting the sub-cloning step into the TOPO-TA vector. Subsequently the nucleotide sequences were then directly ligated into *Eco*RI and *Xho*I digested pSUCTM13ORI generating pBH5 (*gpiB*), pBH11 (*sspM*) and pBH9 (*sspO*) respectively.

The four overexpression constructs were generated by Gibson Assembly [58] of two fragments, respectively. The 2,844 bp backbone fragment, containing a hygromycin resistance cassette, was amplified from pBH12 with the primer pair BH75/76. The nucleotide fragments encoding *gpiB, sspM, sspN* or *sspO* downstream of the P*tef* promoter as well as an ampicillin resistance gene (*bla*) and an *oriT* were amplified from pBH14, pBH18, pBH21 or pBH24 respectively. For amplification, the primer pairs BH72/BH120 (3,476 bp), BH72/BH121 (3,725 bp), BH72/122 (3,525 bp) or BH72/BH123 (3,493 bp) were used, respectively. Assembly of the respective fragments generated the plasmids pBH29 (*gpiB*), pBH30 (*sspM*), pBH31 (*sspN*) and pBH32 (*sspO*).

Constructs encoding the genes of interest, C-terminally fused to an 8xHis tag, were generated via Gibson Assembly [58]. Again, each plasmid was assembled from two fragments. The 2,488 bp backbone was amplified from pBH30 using the BH176/BH75 primer pair. The forward primer, BH176, was designed in a way that it added the 8xHis tag followed by a stop codon to the fragment. The *sspM*-, *sspN*- and *sspO*-encoding fragments were amplified from pBH30, pBH31 and pBH32 with the primer pairs BH72/BH177 (3,728 bp), BH72/BH178 (3,528 bp) and BH72/179 (3,496 bp) respectively. The reverse primers BH177/BH178/BH179 were designed in such a way that the stop codon was removed and an overhang to the 8xHis tag was generated. The *gpiB*-containing fragment was generated based on a previously generated construct, pBH34. To prevent the loss of the 8xHis tag due to the addition of the GPI anchor at the omega site [59], the 8xHis tag was added 30 bp upstream of the omega site. To this end, the vector was assembled in two pieces amplified from pBH34: the 2,823 bp fragment amplified by PCR with BH181/BH75 primer pair containing the C-terminus of *gpiB* as well as the hygromycin resistance cassette, and the 3,377 bp fragment amplified by PCR with the BH72/BH180 primer pair encoding an ampicillin resistance gene, an OriT and the *gpiB* N-terminus. The primers BH180 and BH181 generated the 8xHis tag and thereby overhangs with each other. This generated the plasmids pBH56 (*gpiB*), pBH57 (*sspM*), pBH58 (*sspN*) and pBH59 (*sspO*).

For the localization constructs, the plasmid pBV579 (PN4241) encoding mCherry fused to a T-virus nuclear localization signal, was used [60]. The constructs were generated via Gibson Assembly [58] from three different fragments, generating C-terminal translational fusions between the *ssp* of interest and mCherry-NLS separated by a (GGGS)x2 linker to facilitate independent folding of the protein of interest and mCherry. All constructs incorporated the native *ssp* promoter. The 5,060 bp backbone of the vector was amplified by PCR from pBH12 with the BH76/BH78 primer pair. The 834 bp mCherry-NLS fragment was amplified by PCR from pBV579 with the BH79/BH80 primer pair, where the primer BH79 added the linker. The *sspM, sspN* and *sspO* and the corresponding promoter encoding fragments were amplified by PCR from WT genomic DNA using the primer pairs BH88/BH86 (827 bp), BH91/BH89 (977 bp) and BH93/BH94 (798 bp) respectively. Assembly of the fragments generated the plasmids pBH19, pBH22 and pBH25. All plasmids can be found in S2 Table and all primers in S3 Table.

### Transformation of organisms

Chemically competent *E. coli* DH5α were transformed by heat-shock for 1 min at 42°C, followed by a 2 min incubation step on ice. Cells were allowed to recover for 1 h at 37°C in SOC medium and subsequently plated on LB agar containing ampicillin or kanamycin. *S. cerevisiae* YTK12 cells were made competent using the lithium acetate method [61], transformed by heat-shock for 30 min at 40°C [62] and plated on defined CMD tryptophan dropout medium (see above). *A. tumefaciens* GV3101 [63] cells were made competent and transformed by electroporation (25 mF, 2.5 kV, 200 Ω) [64]. Cells were allowed to recover in SOC medium and then plated on LB agar containing kanamycin. *E. festucae* Fl1 protoplasts were generated as previously described [65]. Protoplasts were transformed with up to 5 µg of the construct using the method previously described [66]. Protoplasts were allowed to regenerate overnight on regeneration agar (RG) and overlaid with RG agar containing either hygromycin B or geneticin the next day.

### Protein extraction, protein purification and western blotting

For protein extraction, *E. festucae* strains were grown for at least 3 days in liquid PD at 22°C. Cultures were filtered through a nappy liner, washed, and snap frozen in liquid nitrogen and ground to a fine powder. Depending on the amount of mycelium, 400–800 µl of lysis buffer (50 mM NaH_2_PO_4_, 300 mM NaCl, 10 mM imidazole, 0.1% (v/v) IGEPAL CA360, 0.2 mM PMSF, 10 µl/ml protease inhibitory cocktail (Sigma Aldrich), 5 mM β-mercaptoethanol) was added and samples were vortexed for 10 min. Subsequently, samples were centrifuged at 14,000 g for 15 min at 4°C and the supernatant was stored at -20°C. The growth medium of each strain (50–80 ml) was collected and filtered through a 0.45 µm filter. Samples were concentrated to approx. 2 ml with Vivaspin 20 MWCO 30,000 spin columns (GE Healthcare Life Sciences) according to the manufacturer’s instructions and washed twice with 10 ml lysis buffer. Samples were stored at -20°C.

The protein purification was performed with Ni-NTA spin columns (Qiagen) according to the manufacturer’s instructions with the exception of the elution buffer (50 mM NaH_2_PO_4_, 300 mM NaCl, 10–500 mM imidazole, 0.1% (v/v) IGEPAL CA360), 0.2 mM PMSF, 10 µl/ml protease inhibitory cocktail (Sigma Aldrich)). Samples were loaded onto 15% (w/v) SDS polyacrylamide gels, electrophoresed at 100 V for 1.5 h then electrophoretically transferred at 30 V for 1 h to PMSF membranes (Roche). The primary antibody, rabbit α-His (Abcam, ab9108), was used in a 1:2,000 dilution and the secondary goat α-rabbit-HRP (Abcam, ab6721) in a 1:10,000 dilution. Detection was performed with the Amersham ECL western blotting detection reagent (GE Healthcare Life Sciences).

For the verification of Ssp and GpiB production in *N. benthamiana*, total protein was extracted from the infiltrated area. First the tissue was snap frozen, then ground to a fine powder in liquid nitrogen and subsequently resuspended in an equal volume of GTEN buffer (10% glycerol, 100 mM Tris pH 7.5, 1 mM EDTA, 150 mM NaCl, 10 mM DTT, 0.2% IGEPAL CA360, 10 µl/ml protease inhibitory cocktail (Sigma Aldrich), 1% PVP). Afterwards, samples were centrifuged at 14,000 rpm for 15 min at 4°C and the supernatant was stored at -20°C. Samples were loaded onto a 15% SDS polyacrylamide gel and transferred onto PMSF membranes (Roche) as described above. The primary antibody, a mouse α-FLAG antibody (Sigma, F6165), was used in a 1:2,000 dilution followed by the secondary goat α-mouse–HRP (Santa Cruz Biotechnology, Dallas, TX, USA) antibody in a 1:1,000 dilution.

### RNA extraction and qPCR

For the determination of the gene copy number, high quality gDNA was extracted as described above and qPCR was performed with SYBR Green (Invitrogen) on a LightCycler 480 System (Roche) as per the manufacturer’s instructions. Each sample was analysed with two technical replicates. The single copy genes *hepA* (EfM3.043690) and *pacC* (EfM3.009480) were used as reference genes and the copy number was calculated relative to WT as described previously [67].

For the determination of relative expression levels, RNA was extracted from freeze-dried mycelium using TRIzol reagent (Invitrogen). Subsequently, cDNA synthesis was performed with the QuantiTect Reverse Transcriptase kit (Quiagen). The qRT-PCR was performed as described above. Each sample was analysed with two technical replicates. The genes coding for translation elongation factor 2 (EF-2, EfM3.021210) and 40S ribosomal protein S22 (S22, EfM3.016650) were chosen as reference genes as previously described [68]. The expression level was calculated relative to WT as described previously [67]. All primers used for qPCR and qRT-PCR can be found in the S3 Table.

### Analysis of transmission of ***E. festucae* mutants into *L. perenne* seeds**

Twenty endophyte free seedlings of *L. perenne* cv. Samson were inoculated with three independent deletion strains of *gpiB* (T111, T133, T148), *sspM* (T52, T99, T163), *sspN* (T10, T30, T52) and *sspO* (T78, T195, T210) [56, 69]. Twenty seeds were inoculated with PD agar as mock control. Plants were grown in root trainers in the greenhouse for 7 weeks at 20°C with a photoperiod of 16 h of light and tested for presence of the endophyte by immunoblotting [70]. Five immunoblot positive plants of each strain were transferred to new pots 10 weeks post-infection and grown in the greenhouse for additional two weeks. For vernalisation, plants were transferred to a growth cabinet and cultivated for 7 weeks at 6°C with a short-day photoperiod of 8 h light and 16 h dark. Starting from four weeks until ten weeks after vernalisation, plants produced inflorescences. Ovaries were collected, stored in 95% EtOH and stained [69] when pollen was visible and ovaries mature. Stained ovaries were stored in 70% glycerol until microscopic observation. Seeds were collected, dried and stored in the fridge until further use. For microscopy and immunoblotting of seeds, the seeds were surface sterilised [56, 69], germinated on wet filter paper overnight and cross sections used for immunoblotting.

### Microscopy

For microscopy of *E. festucae* culture growth, a small piece of a fresh colony was inoculated on the edge of a glass slide covered with a thin layer of 1.5% H_2_O agar placed on an agar plate with 3% H_2_O agar. Strains were incubated for 6 days before examination with an Olympus IX71 inverted fluorescence microscope using the filter sets for DIC and CFW/DAPI. For staining of the cell wall, a drop of a 3 mg/ml of Calcoflour White was added to the sample just before microscopy.

For the examination of growth and morphology of hyphae *in planta*, pseudostem tissue was stained with aniline blue diammonium salt (Sigma) and Wheat Germ Agglutinin conjugated to AlexaFluor^®^488 (WGA-AF488, Molecular Probes/Invitrogen). First, infected tissue was incubated in 100% EtOH overnight at 4°C followed by an incubation in 10% KOH overnight at 4°C. Samples were washed at least 3 times with PBS (pH 7.4) before incubation in the staining solution (0.02% aniline blue, 10 ng/ml WGA-AF488, 0.02% Tween^®^-20 (Invitrogen) in PBS (pH 7.4)) for 10 min. Samples were vacuum-infiltrated with the staining solution for 30 min and then stored at 4°C until analysis. Examination of these samples was performed with a Leica SP5 DM6000B confocal microscope (488 nm argon and 561 nm DPSS laser, 40× oil immersion objective, NA= 1.3) (Leica Microsystems). For TEM pseudostem samples were fixed with 3% glutaraldehyde and 2% formaldehyde in 0.1 M phosphate buffer, pH 7.2 for 1 h, as described previously [71]. The fixed samples were examined with a Philips CM10 TEM and the images were acquired using a SIS Morada digital camera. Sections of the resin-embedded samples were also stained with 0.05% toluidine blue in phosphate buffer and heat-fixed at approx. 100°C for 10 s. These samples were examined with a Zeiss Axiophot Microscope with Differential Interference Contrast (DIC) Optics and Colour CCD camera.

For the *in planta* localization of the Ssp-mCherry-NLS fusion proteins, un-fixed pseudostem samples were examined with a Leica SP5 DM6000B confocal microscope (488 nm argon and 561 nm DPSS laser, 40× oil immersion objective, NA= 1.3).

### Preparation of binary vectors, agroinfiltration and suppression assay

All binary backbone vectors for transformation into *Agrobacterium* are listed in S2 Table. The plasmids pBH44-47 were generated via Golden Gate cloning using pICH86988 as the backbone. The nucleotide sequences of *gpiB, sspM, sspN and sspO* were amplified by PCR without their native secretion signal from WT *E. festucae* cDNA with primer pairs adding additional base pairs for the necessary overhang and a *Bsa*I restriction site (*gpiB*: BH165/BH166, *sspM*: BH167/BH168, *sspN*: BH170/BH171, *sspO*: BH172/BH173) as described. The *N. tabacum* secretion signal PR1α was obtained from pUC19B (S. Marillonet, pers.comm.). The digestion and ligation were conducted either in one step or in two separate steps. The one-step reaction was performed in a total volume of 20 µl with 1 µl T4 ligase (NEB) and 1 µl BSA (NEB) and equimolar amounts of all three inserts (approx. 50 ng total). The reaction was performed in a Thermocycler (Eppendorf) with the following settings: 25 cycles of 37°C for 3 s and 16°C for 4 s, followed by 5 s inactivation at 50°C and 80°C each. For the two-step reaction, the digest was performed in a volume of 10 µl with 1 µl BSA (NEB) and equimolar amounts of each insert. The reaction was performed at 37°C for 3 h. Subsequently the reaction was inactivated by heating to 50°C. For the ligation, the volume was topped up to 20 µl per reaction including 1 µl T4 ligase (NEB). The reaction was incubated at 16°C in a Thermocycler (Eppendorf) overnight followed by an 80°C inactivation step. A diagnostic restriction digest and DNA sequencing confirmed the sequence of all vectors. Electro-competent *A. tumefaciens* GV3101 cells were generated and transformed as described. Cells were allowed to recover in SOC medium and subsequently plated on LB agar containing kanamycin. To perform the infiltrations, fresh strains were inoculated in 3 ml LB medium with kanamycin and incubated at 30°C overnight. The next day, cultures were spun down and resuspended in 1 ml infiltration buffer (10 mM MgCl_2_, 10 mM MES-KOH, 100 µM acetosyringone). These cultures were adjusted to an OD_600_ of 0.4 in the necessary amount of infiltration buffer (usually 5–10 ml) and incubated at room temperature for at least 3 h before infiltration. To confirm the expression of each of the constructs, total protein was extracted from infiltrated *N. benthamiana* tissue and a western blot was performed.

For HR suppression assays *A. tumefaciens* strains carrying plasmids encoding the potential suppression gene pBH44 (*gpiB*), pBH45 (*sspM*), pBH46 (*sspN*), pBH47 (*sspO*), pBIN-Plus-Avr3a (suppression control), pICH86966-N-3xFlag-GFP (control), pBC302-3-R3a (HR control) were infiltrated 24h before infiltration with pBC302-3-INF1 and pBIN-Plus-Avr3a (HR control).

## Results

### Candidate effectors of *E. festucae*

The recent availability of the complete telomere-to-telomere genome sequence of *E. festucae* strain Fl1, which is 35 Mb in size and encodes 8,465 predicted genes on seven chromosomes [26], provided us with the opportunity to study the organization, structure and genome context of genes encoding candidate effector proteins. First, we established which of the 8,465 predicted genes likely encode secreted proteins. For this purpose, we defined a secreted protein as any protein with a predicted amino N-terminal secretion signal, as determined using the SignalP server, that targets the protein outside of the fungal plasma membrane [44]. This definition does not exclude those proteins with a predicted transmembrane domain or GPI anchor modification site. In total, 682 genes (8.1%) were predicted to encode a secreted protein. We then analysed which of the 682 predicted secreted proteins encode candidate effectors using the EffectorP server, which predicts effectors using a machine learning approach [41]. In addition, we applied a size cutoff of ≤200 amino acids, as most fungal effectors fall within this range [72]. Out of 175 predicted secreted proteins meeting this length requirement, 141 were predicted to be candidate effectors by EffectorP. For subsequent analyses the 8,465 predicted proteins were subdivided into ‘effector candidates’ (n=141), ‘secreted’, comprising all putatively secreted proteins that are not classified as effector candidates (n= 541), and ‘non-secreted’ made up of all putatively non-secreted proteins (n= 7,783) (S4 Table).

To gain further insight into the possible functions of the 141 effector candidates, and to determine which of these predicted proteins have homologs across the *Sordariomycetes*, a BLASTp analysis was performed against fungal reference proteins at NCBI. Of the 141 effector candidates, 83 were found to have sequence homology to proteins present in the NCBI database (E-value cut off of ≤1e-10), but 56 of these were to uncharacterized or hypothetical proteins. Among the ‘hits’ to characterized proteins were ten cell wall-associated proteins, including hydrophobins and carbohydrate-binding proteins, two toxins and other small secreted proteins (S4 Table). The majority of ‘hits’ were to proteins from species in the Hypocreales (21 *Claviceps spp*., 21 *Metarhizium spp*., 14 *Pochonia spp*., 6 *Moelleriella spp*., 6 *Ustilaginoiea spp*.), with the remaining hits to proteins from other species within the Sordariomycetes. For 58 of the effector candidates, no homologs could be identified (E-value>1e-10), but analysis against a custom made Clavicipitaceae endophyte database (epichloe.massey.ac.nz)[47] found potential homologs for 20 of these proteins. The remaining 38 Ssps are apparently unique to *Epichloë spp.* (S4 Table).

Given many fungal effectors are cysteine-rich, a property that enables the formation of disulfide bonds and protein folding to maintain structural integrity in the protease-rich apoplast [73, 74], the frequency of cysteine residues in the effector candidates was compared with that of the secreted and non-secreted categories. As expected the candidate effectors were significantly more cysteine-rich (3.6% of residues) than either secreted (frequency= 1.7%, *P*= <1e-18) or non-secreted (frequency= 1.4%, *P* < 1e-18) (S1 File).

In smut fungi, secreted genes are predominantly arranged in clusters within the genome and are highly upregulated during tumor formation [75, 76]. We therefore tested whether the genes encoding the *E. festucae* effector candidates were clustered by mapping them to each of the seven chromosomes (Fig 1A). From this low-resolution analysis, it appears that the genes encoding effector candidates of *E. festucae* are evenly distributed throughout the genome. This conclusion was supported by an analysis comparing the within category gene distances, which concluded that genes encoding effector candidates are not clustered in the *E. festucae* genome (Fig 1A). Of all the genes encoding candidate effectors, only eight (four gene pairs) were found within 1 kb of each other. In fact, the mean distance between candidate effector genes was not significantly different from that expected from a random subset of the 141 genes (*P*= 0.236).

**Fig 1.**
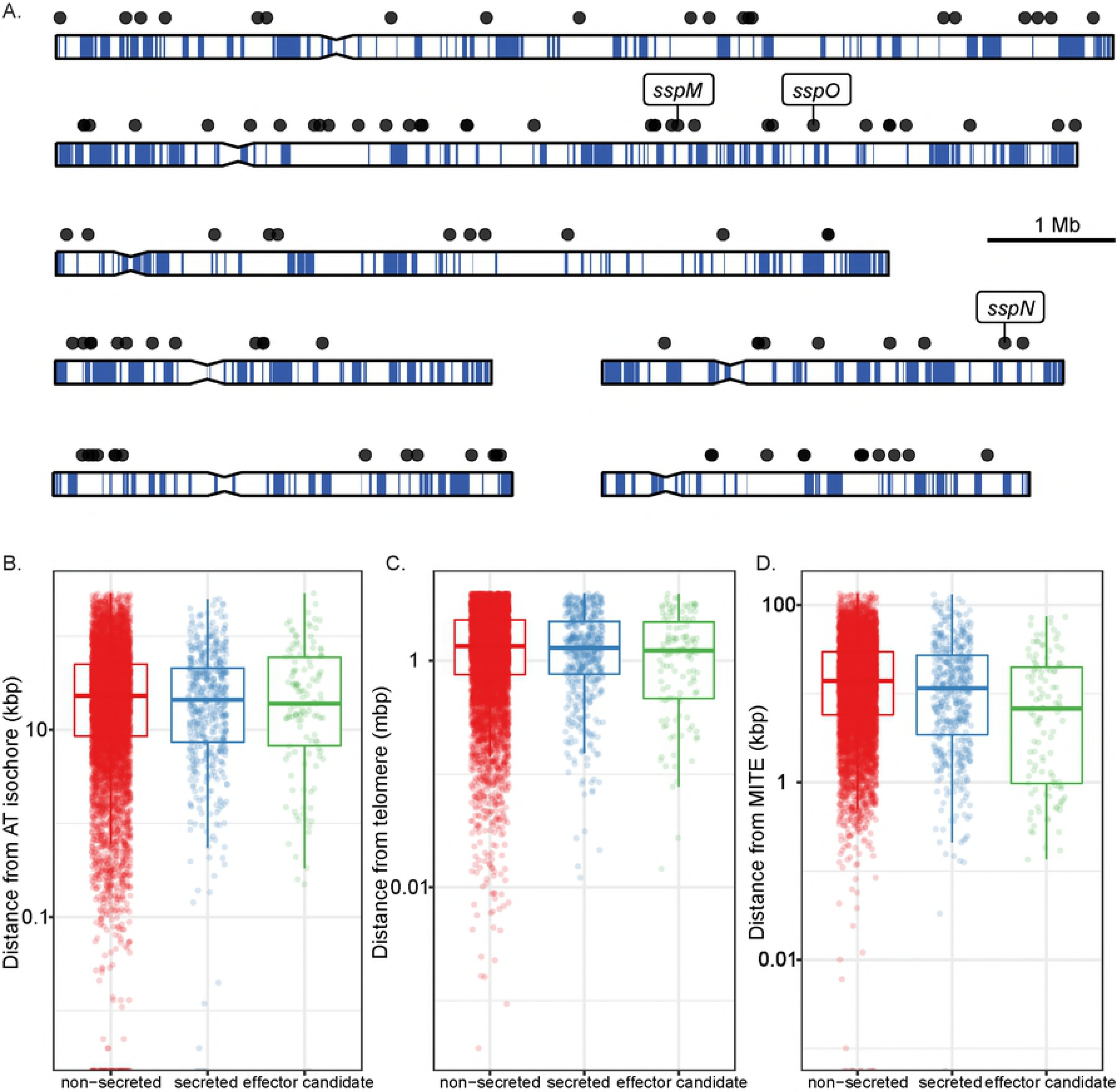
Physical properties of genes encoding effector candidates in *E. festucae*. (A) Location of genes encoding effector candidates (black circles) on the seven chromosomes of *E. festucae* strain Fl1. The blue line indicates AT richness in a 1,000 bp window. (B) Distance of each *E. festucae* gene from AT-rich regions (isochores). (C) Distance of each *E. festucae* gene from telomeres. (D) Distance of each *E. festucae* gene from MITES. Genes are divided into the categories ‘non-secreted’ (n= 7783), ‘secreted’ (n= 541) and ‘effector candidates’ (n= 141).

*E. festucae* is one of several filamentous fungi that has a genome comprised of distinct gene-poor AT isochores and gene-rich GC isochores [21, 26]. Because the AT-rich regions are subject to an unusually high mutation rate [77-80] they are frequently referred to as ‘birthplaces’ for new genes such as those encoding effectors and secondary metabolites [81]. In *Leptosphaeria maculans* 20% of all genes in AT isochores encode small secreted proteins (SSPs, effector candidates), whereas they make up just 4.2% of all genes in GC isochores [21]. In contrast to *L. maculans*, there was no enrichment of genes encoding effectors in or near to AT isochores of the *E. festucae* genome (*P*= 0.369) (Fig 1B). Similarly, there was no enrichment of genes in the secreted category in AT isochores (*P*= 0.812) (Fig 1B). Indeed, of the 548 genes located in or within 1,000 bp of an AT isochore, only 1.5% and 4.7% encoded a protein from the candidate effector and secreted categories, respectively.

Given some fungal effector genes, such as the *AVR-Pita* gene from *Magnaporthe oryzae* [82], are found in close proximity to telomeres, we took advantage of the telomere-to-telomere *E. festucae* genome sequence to determine if genes encoding candidate effectors are located close to telomeres. These regions of the genome are also subject to higher mutation rates than the remainder of the genome and potential sites for the evolution of environment adaptation genes such as those encoding effectors [83, 84]. However, genes encoding effector candidates were not significantly closer to telomeres than other genes present in the *E. festucae* genome (*P*< 0.787) (Fig 1C).

Transposable elements are increasingly seen as important regulators of gene expression [85, 86], and miniature inverted-repeat transposable elements (MITES) have been postulated to play a role in the expression of *E. festucae* genes *in planta* [24, 26]. For this reason, the distance between genes encoding effector candidates and MITES was analyzed in the *E. festucae* genome. This analysis revealed that genes encoding effector candidates are more likely to contain a MITE within 2 kb of their transcription start site than non-secreted proteins (*P*= 3.56e-18, Fig 1D). Genes encoding other secreted proteins were also more likely to contain a MITE immediately upstream of their transcription start site than non-secreted proteins (*P*= 1.66e-7). No particular MITE family is over-represented within 2 kb of these genes.

Apart from the physical organization of effector genes within fungal genomes, the expression of these genes often follows distinct host colonization patterns and are generally more highly expressed *in planta* than in axenic culture [72, 87, 88]. We therefore analysed the transcript expression profiles of the three different gene categories between *E. festucae* grown *in planta* and in axenic culture. Genes encoding effector candidates were significantly more highly expressed *in planta* (mean log_2_ fold difference= 1.20, *P*< 2e-16) (Fig 2A). Other secreted proteins were also moderately more highly expressed *in planta* (mean log_2_ fold difference= 0.20, *P*= 0.004) (Fig 2A).

**Fig 2.**
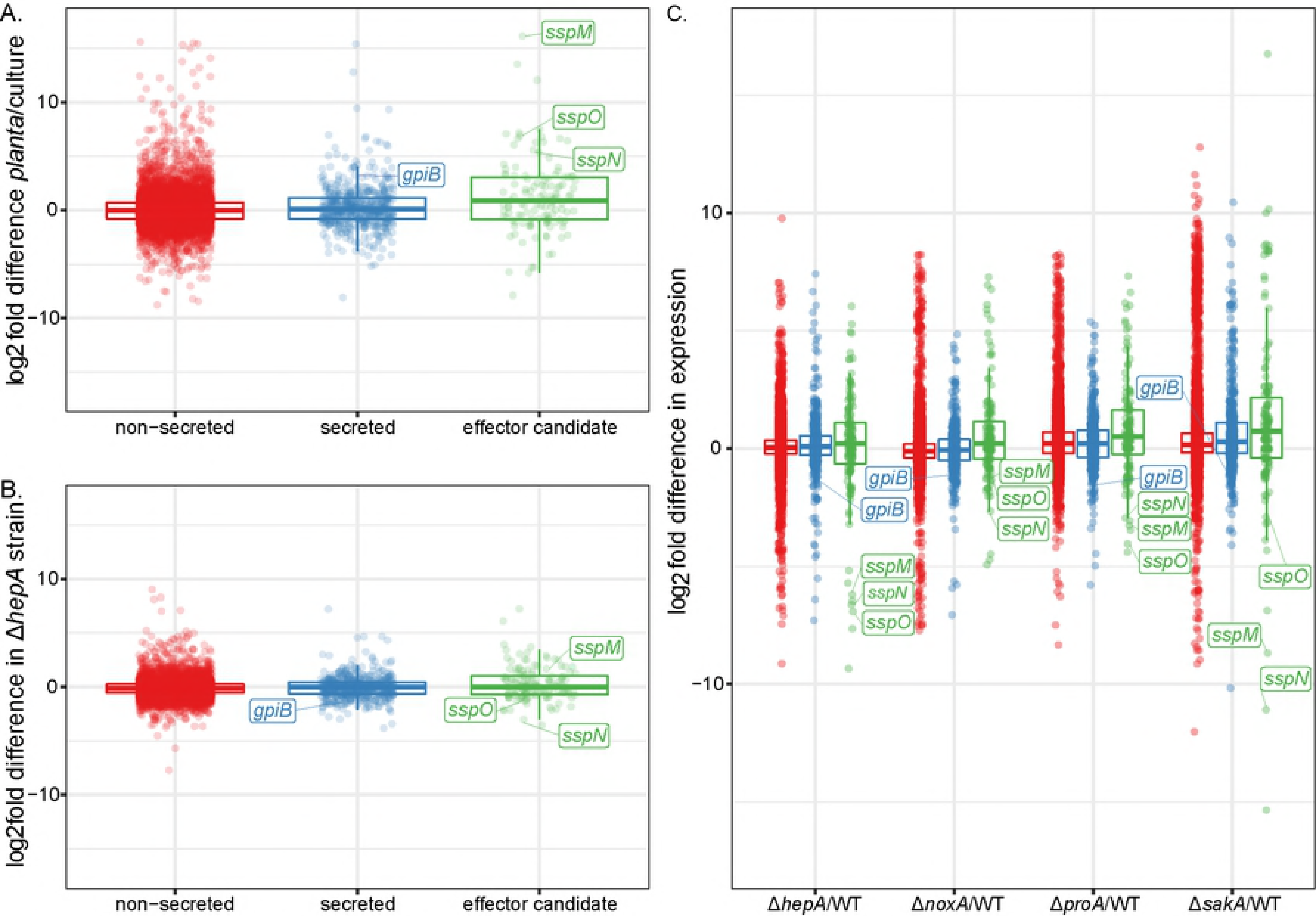
Analysis of gene expression in WT *E. festucae* and symbiotic mutants of *E. festucae* grown. (A) Log_2_ fold difference in gene expression between WT *E. festucae* grown *in planta* and in culture. (B) Log_2_ fold difference in gene expression between WT and a *hepA* deletion strain grown in culture. (C) Log_2_ fold difference in gene expression between WT and Δ*noxA,* Δ*proA,* Δ*sakA* and Δ*hepA* strains grown *in planta*. Number of genes in each category: ‘non-secreted’, n= 7783; ‘secreted’, n= 541; and ‘effector candidates’, n= 141.

Given genes encoding effector candidates are preferentially silenced in axenic culture compared to *in planta* we examined whether this might be due to the chromatin state as shown for *L. maculans* where RNAi-induced silencing of DIM-5, a histone H3 lysine 9 trimethyl transferase, and heterochromatin protein 1 (HP1), led to de-repression of effector gene expression in axenic culture [89]. We therefore compared expression of *E. festucae* genes encoding effector candidates in a *hepA* (homolog of *HP1*) mutant versus wild type. We found candidate effector genes were moderately more highly expressed in the *hepA* mutant (mean log_2_ fold difference= 0.310, P= 4.13e-9) (Fig 2B). For some genes, this change in gene expression was substantial. Eleven percent of putative effectors showed log_2_ fold difference in expression greater than 2, a significantly higher proportion than observed in larger secreted (1.5% of genes, P= 5e-14) or non-secreted proteins (3% of genes, P= 1e-4)).

Previously we showed that disruption of the usually asymptomatic interaction between *L. perenne* and *E. festucae* by infecting the host with symbiotic mutants of *E. festucae* results in major changes in the fungal transcriptome [47]. We therefore tested whether effector candidate gene expression changed in the Δ*sakA*, Δ*noxA,* Δ*proA* and Δ*hepA* symbiotic mutants using previously generated transcriptome data sets [47](Tetsuya & Scott, personal communication). Generally, genes encoding effector candidates were upregulated in each of the dysfunctional interactions (mean difference of 0.411 log2 fold units, P = 5.28 x 10-8) (Fig 2C). A closer investigation showed that ten genes encoding effector candidates SspB-SspK were consistently upregulated across all four different mutant interactions and three genes encoding effector candidates SspM-SspO were consistently down regulated [47](Tetsuya & Scott, personal communication).

In summary, the genes encoding effector candidates from *E. festucae* do not show a distinct distribution pattern as has been described for some other fungi, but seem to be evenly dispersed throughout the genome. These genes do not cluster, and are not preferentially located in AT-rich isochores or telomeres. However, these genes are enriched with respect to their proximity to MITES. As found for many other fungi, a significant number of genes encoding effector candidates from *E. festucae* are preferentially upregulated *in planta*. Furthermore, many of these genes are upregulated in symbiotic mutants of *E. festucae* that disrupt the restrictive pattern of growth observed for WT *E. festucae*.

### Sequence analysis of downregulated effector candidates

To gain further insight into the role of *E. festucae* effectors in the symbiotic interaction with *L. perenne*, the three genes described above (*sspM*, EfM3.016770; *sspN*, EfM3.062700; and *sspO*, EfM3.014350) that were found to be downregulated in all four symbiotic mutant interactions, were selected for genetic analysis [47](Tetsuya & Scott, personal communication). The fact that they were among the top 100 genes upregulated *in planta* compared to axenic culture was an additional criterium for selecting these three genes (Fig 2A, S4 Table). A fourth gene, *gpiB* (EfM3.018200), encoding a putative GPI-anchored protein, was also selected for functional analysis because it had an identical expression profile to *sspM, sspN* and *sspO* [47], and because GPI-anchored proteins are known to have important roles in fungal virulence [42, 43].

cDNA sequencing confirmed that the proposed gene models for each of these four genes were correct. In the case of *sspN*, where two alternatively spliced forms were proposed the second of these (mRNA-2) corresponded to the cDNA sequence. The *gpiB* gene (EfM3.018200) is predicted to encode a protein of 126 aa with an unmodified MW of 12.61 kDa (8.7 kDa without secretion signal and C-terminus beyond the omega cleavage site), while the *sspM* gene (EfM3.016770) is predicted to encode a protein of 177 aa with an unmodified MW of 18.87 kDa (17.1 kDa without secretion signal). The protein encoded by *sspN* (EfM3.062700) is predicted to be 127 aa in length with an unmodified MW of 14.98 kDa (13.19 kDa without secretion signal), while the protein encoded by *sspO* (EfM3.014350) is predicted to be 104 aa in length with an unmodified MW of 11.11 kDa (9.4 kDa without secretion signal) (Fig 3).

Analysis of the selected protein sequences with InterproScan [90], SMART [91], T-Reks [92] and HHPRED [93] did not identify any conserved domains with known function, but did predict a number of different structural features and secondary modifications (Fig 3). GpiB contains a predicted *N*-glycosylation motif as well as a putative C-terminal GPI anchor. The SspM protein contains a possible pre-pro domain ending with a classical LxKR kexin cleavage site motif and a predicted *N*-glycosylation site. SspN is a repeat-containing protein, consisting of 7.5 almost perfect repeats of 15 aa. Like SspM, it also contains a putative kexin cleavage site. Analysis of the SspO sequence did not reveal any obvious structural features. Both SspM and SspO are rich in cysteines; SspO containing eight and SspM containing six downstream of the cleavage site, whereas SspN and GpiB are cysteine-free (Fig 3).

**Fig 3.**
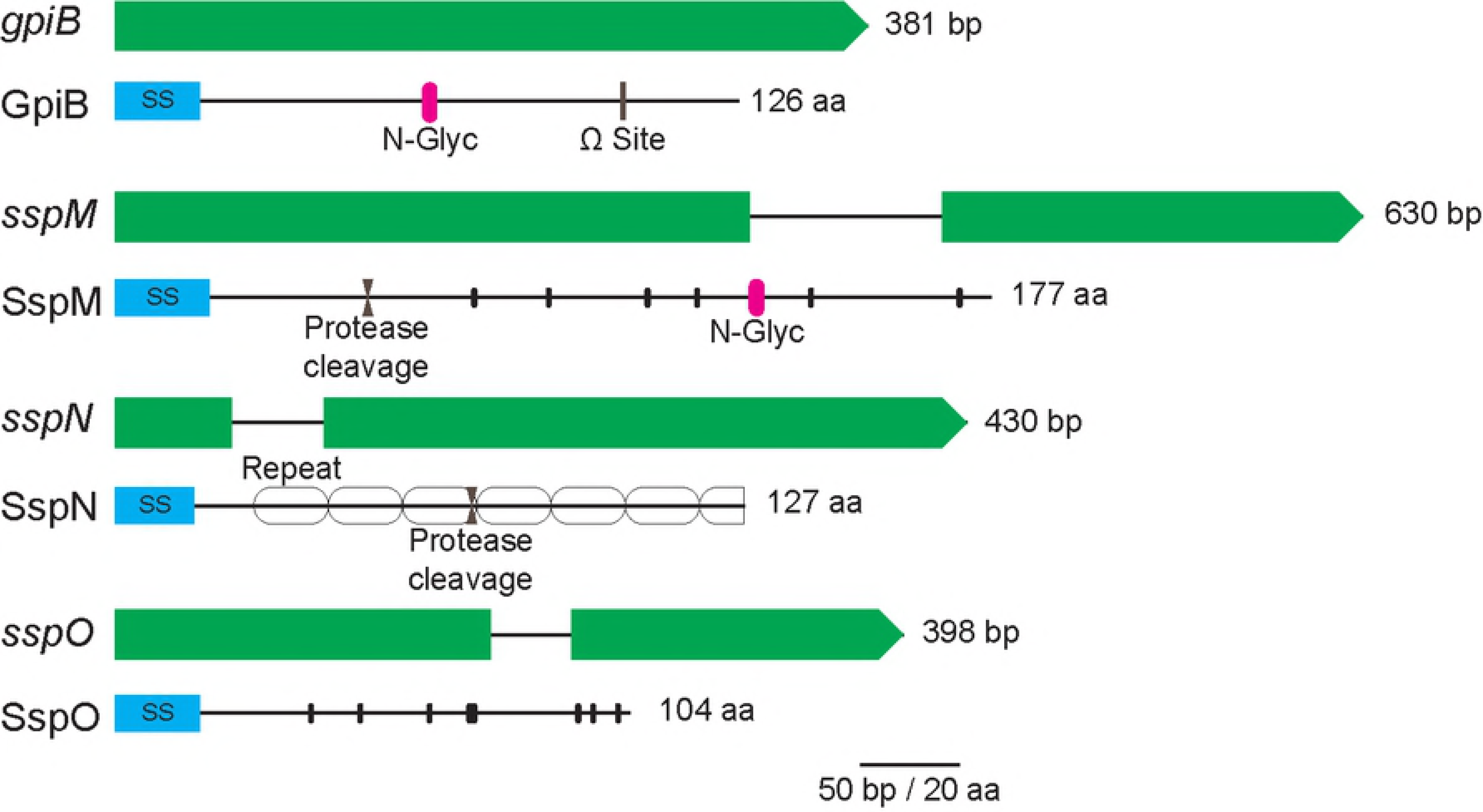
Schematic representation of the small secreted proteins SspM, SspN, SspO (effector candidates) and the GPI-anchored protein GpiB and their corresponding genes. GpiB (EfM3.018200) is encoded by a 381 bp gene that lacks introns, and is predicted to encode a protein of 126 aa. The gene encoding the predicted 177 aa SspM (EfM3.016770) is 630 bp in size and contains one intron. SspN (EfM3.062700) is encoded by a gene of 430 bp, with one intron, and encodes a predicted protein of 127 aa. SspO (EfM3.014350) is encoded by a 398 bp gene, containing one intron, and is predicted to be encode a protein of 104 aa. N-terminal secretion signals (SS) are indicated by blue boxes and post-translational modifications and structural motifs by pink bars (*N*-glycosylation), grey bars (GPI anchor modification site), facing arrows (protease cleavage site), hollow ovals (repeats) and small black bars (cysteine residues).

A BLASTn analysis of the four selected genes revealed a patchy distribution within the *Epichloë* species, with only *gpiB* present in all analyzed genomes available on the Kentucky Endophyte Database [94]. The observed distribution patterns did not correlate with specific hosts or the sexual phenotype of the fungus (S5 Table). Alignment of the protein sequences of interest with the corresponding homologs in other *Epichloë* species showed that GpiB and SspO are highly conserved (S1A and S1D Figs). Interestingly, SspM was highly conserved from the N-terminus to the proposed kexin cleavage site and less conserved beyond this site (S1B Fig). The repeat structure of SspN was also conserved, with homologs mainly differing in their number of repeats (S1C Fig). A BLASTp analysis identified homologs of Ssps and GpiB in species outside of *Epichloë* (S4 Table); however, none of these are functionally characterized.

### GpiB, SspM, SspN and SspO are secreted

To experimentally verify secretion of the three selected candidate effectors and the GPI anchored protein, GpiB, a complementation assay was performed in *S. cerevisiae*. The nucleotide sequence encoding the secretion signal predicted by SignalP was ligated in frame to the N-terminus of the *S. cerevisiae* invertase *SUC2* encoded on the pSUC2T7M13ORI vector [52]. These constructs were then transformed into the *SUC2*-negative yeast strain YTK12 and tested for growth on CMDRAA medium, which contains raffinose as the sole carbon source. All four test strains expressing the secretion signal-Suc2 fusion protein grew on this medium whereas the strain expressing Suc2 without a secretion signal and the untransformed YTK12 did not (Fig 4). These results indicate that the signal peptides of SspM, SspN, SspO and GpiB are sufficient to mediate secretion in *S. cerevisiae*.

**Fig 4.**
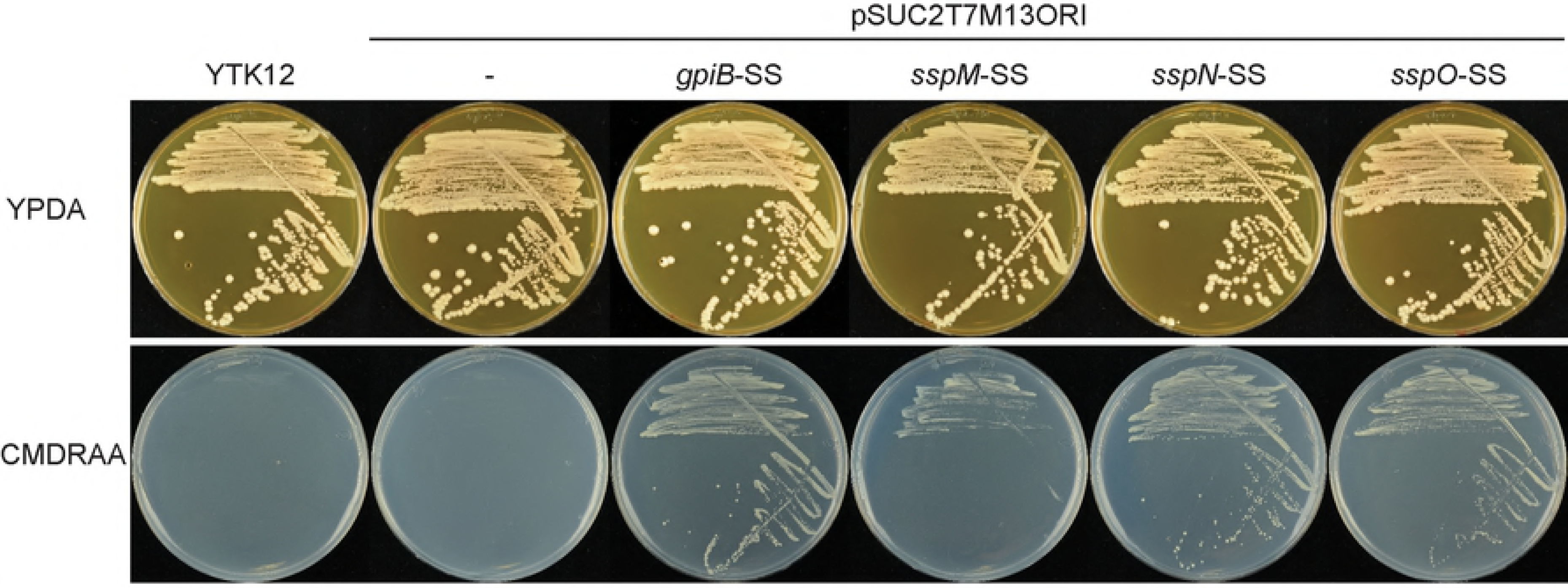
Analysis of the GpiB, SspM, SspN and SspO secretion signals in a *S. cerevisiae* complementation assay. YTK12 strains transformed with pSUC2T7M13ORI encoding *SUC2* transcriptionally fused to *gpiB, sspM, sspN* or *sspO* secretion signal nucleotide sequence, as well as the empty vector and YTK12-only controls. Only strains expressing the secretion signal-Suc2 fusion proteins were able to grow on defined medium with raffinose as carbon source (CMDRAA). SS, secretion signal. All strains grew on the control plates containing YPDA media.

### Analysis of secretion and processing of GpiB, SspM, SspN and SspO in *E. festucae*

As discussed above, many effectors are post-translationally processed (e.g. by proteolytic cleavage) or modified (e.g. by glycosylation) to their final form. To verify the secretion of these proteins in *E. festucae*, and to check if they are modified (Fig 3), His-tagged fusions of GpiB, SspM, SspN and SspO were purified from extracts of *E. festucae* mycelia (intracellular form), as well as from the liquid culture medium (extracellular form), separated by PAGE and analyzed by western blotting (Fig 5). All four proteins were present in extracts from fungal mycelia. The 13 kDa band for GpiB is larger than the predicted His-tagged size of 9.8 kDa, possibly due to the addition of the GPI anchor and/or modification by *N*-glycosylation. A band corresponding to SspN was detected at approx. 14 kDa, which agrees with the predicted His-tagged size of 14.31 kDa for the unprocessed protein, indicating that under the growth conditions used, this protein is not cleaved, even though it has a predicted kexin cleavage site. The signal for SspO was very faint, and slightly larger (12 kDa) than the predicted His-tagged size of 10.5 kDa for this protein. In contrast, two bands of 20 and 16 kDa were detected for SspM, with the band of higher molecular weight being very faint. SspM has a predicted His tagged size of 18.22 kDa in the unprocessed form (with its predicted pro-domain) and 12.7 in the cleaved form (without its predicted pro-domain). Taking into account a probable increase in size due to the predicted *N*-glycosylation, the sizes observed for SspM agrees with the predicted sizes of the kexin-unprocessed and processed forms, respectively. The presence of additional bands on the western is probably the result of non-specific binding to other proteins present in the crude extract separated on these gels.

**Fig 5.**
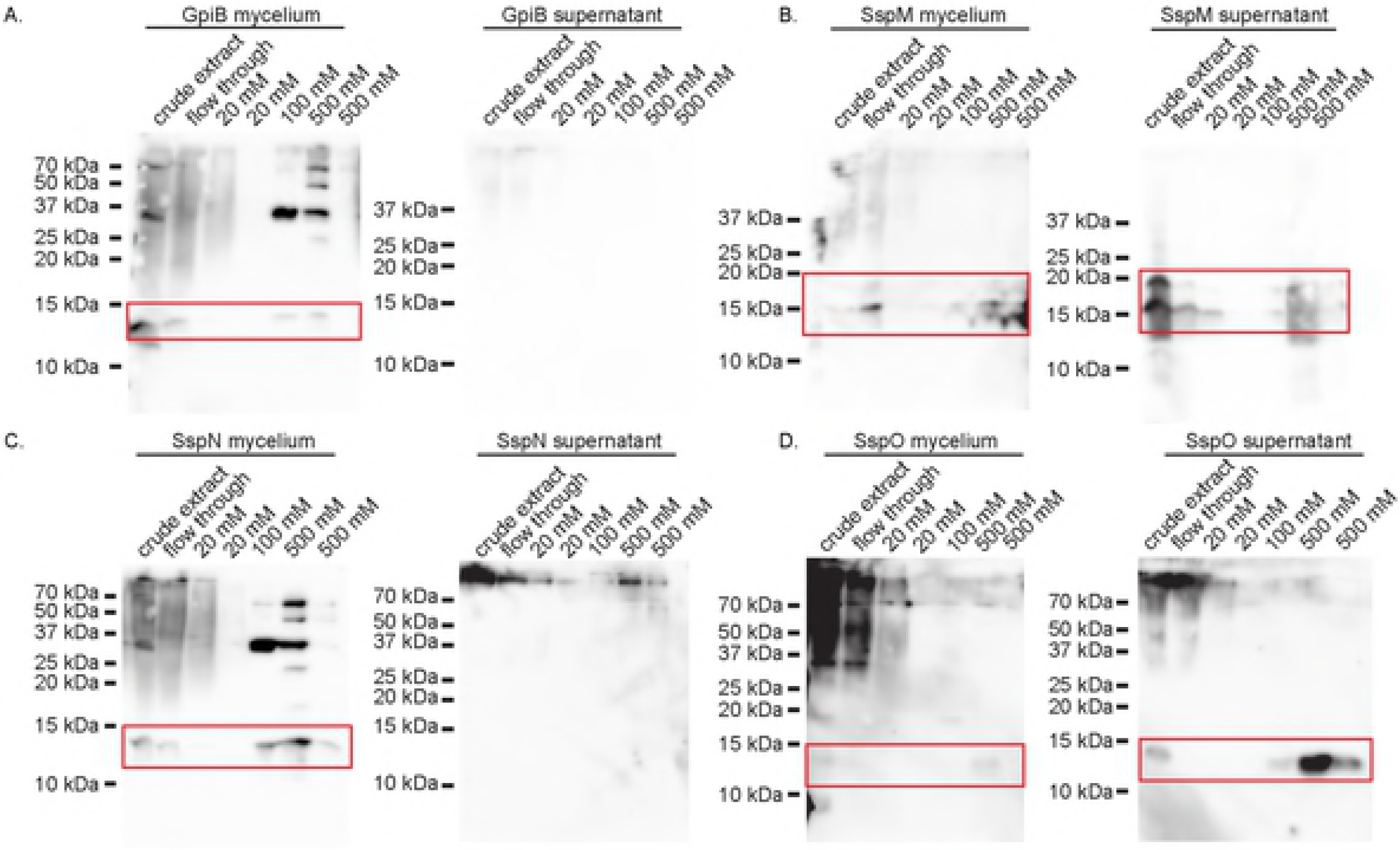
Western blot analysis of GpiB-His, SspM-His, SspN-His and SspO-His extracted from *E. festucae* mycelia and culture medium. Genes of interest (*gpiB, sspM, sspN* and *sspO*) were translationally fused to an 8x His tag at the C-terminus and constructs were placed under the control of the P*tef* overexpression promoter. Total protein was extracted from fungal mycelia and the corresponding growth medium, and His-tagged proteins purified on Ni-NTA spin columns eluted with increasing concentrations of imidazole. Samples were separated on a 15% SDS PAGE gel, transferred to membranes and probed with an anti-His antibody. SspM-His was extracted from *E. festucae* strains grown in medium buffered to pH 6.5 with 50 mM HEPES. Predicted sizes of proteins were calculated on the basis that each protein lacked the secretion signal and had the 8x His-tag.

Analysis of the culture medium identified bands for SspO and SspM, but the latter was only detected when the culture medium was buffered to pH 6.5 with 50 mM HEPES. The bands detected for SspO and SspM were the same size as those seen in the mycelial fraction, with the latter again present as a double band. These results suggest that SspO does not contain any secondary modifications and is secreted into the culture medium, whereas SspM is processed at the predicted kexin site. The absence of bands for GpiB and SspN in the culture medium suggest that these proteins are either unstable in the culture medium or remain attached to the fungal hyphae following secretion.

### Functional analysis of *gpiB, sspM, sspN and sspO*

To investigate whether *sspM, sspN, sspO* or *gpiB* are necessary for establishing a mutualistic symbiotic interaction, given their expression is downregulated in three different symbiotic mutants [47], each gene was individually deleted in the WT background using a gene replacement approach. Replacement constructs, pBH1–4, were prepared, and PCR-amplified linear fragments of each were transformed into WT protoplasts. Putative gene replacements were identified by PCR screening approx. 200 independent Hyg^R^ transformants with primers that flank the site of *hph* integration. Additional PCR screening with primer sets that amplify the left and right flanks of each of the insertion sites was used to confirm the transformants selected had gene replacements. Southern blot analysis of genomic DNA extracted from these strains identified three independent deletion strains for *gpiB* (T111, T133, T148), *sspM* (T52, T99, T163*) sspO* (T78, T195, T210), and *sspN* replacement (T10, T30, T52) (S2 Fig).

To determine the phenotype of the deletion strains in culture, the whole colony morphology of each independent deletion strain was compared to that of WT, which is typically circular with white fluffy mycelium that thins and flattens towards the edges. This comparison revealed that the whole colony morphology of each deletion strain was indistinguishable from WT (Fig 6). Given this result, the hyphal morphology was more closely analysed by microscopy. *E. festucae* WT had smooth hyphal tips with hyphae arranged in bundles that are thicker in the more mature inner zone of the colony. Hyphae of WT had hyphal bridges, which are formed by tip-to-side fusion [40], as well as hyphal coil structures, which are thought to promote conidiophore formation [40]. When stained with Calcofluor white, which preferentially binds to chitin, the cell walls of WT hyphae uniformly fluoresced. Similar to the whole-colony comparison, each of the deletion strains had an indistinguishable phenotype to that of WT, forming hyphal bundles, with occasional hyphal fusion, and coil formation (Fig 6).

**Fig 6.**
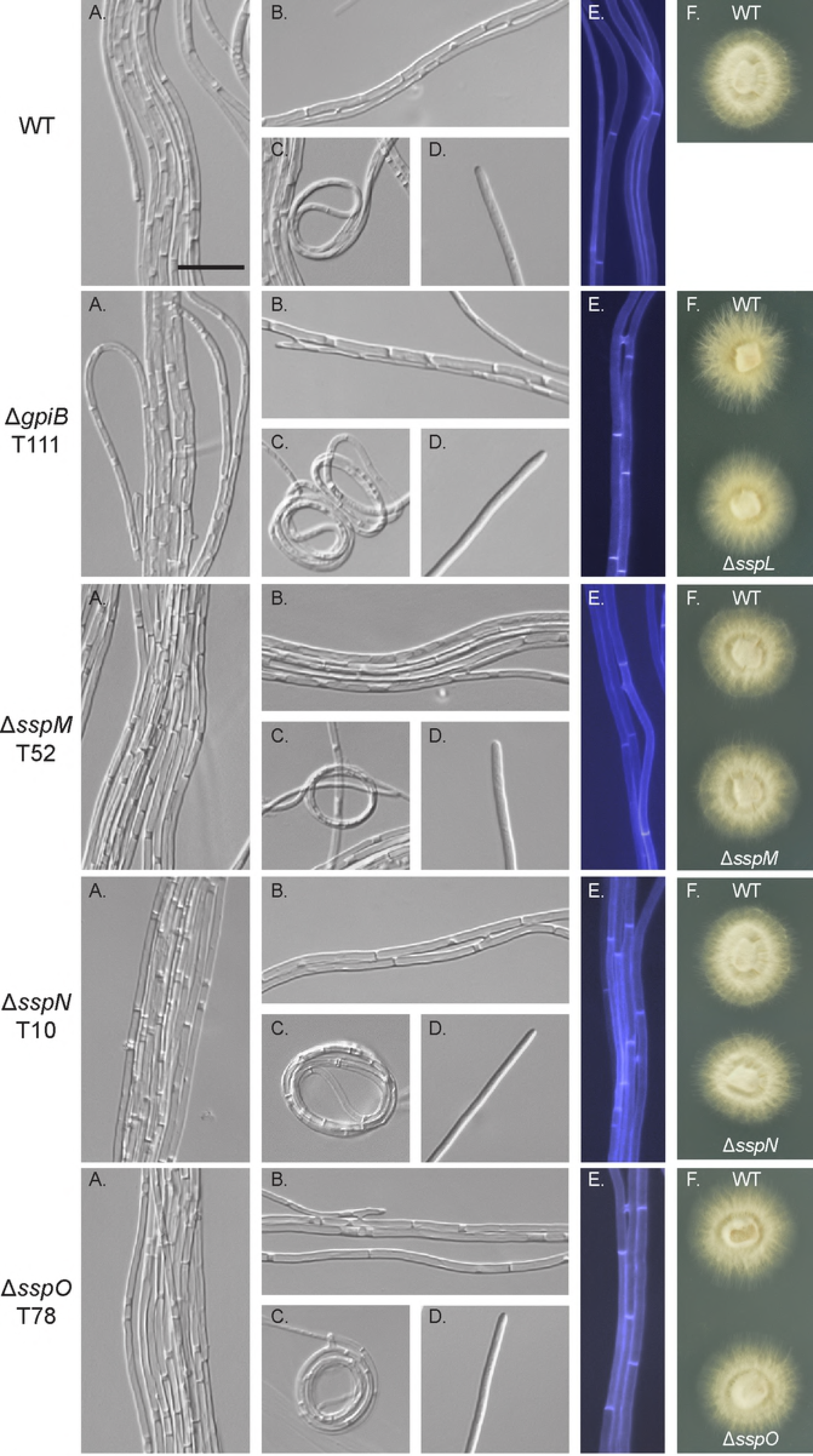
Culture phenotype of WT and deletion mutant strains. (A)-(D) Representative DIC images captured with the inverted microscope of WT and deletion mutant strains grown on 1.5% water agar for 6 days. (A) Hyphal bundle formation. (B) Hyphal fusion points. (C) Hyphal coils. (D) Hyphal tips. (E) Hyphae stained with Calcofluor white to examine cell wall composition. (F) Colony morphology of WT compared to the deletion strains grown on 2.4% PDA for 7 days. Bar: 20 µm.

To determine the host interaction phenotype, strains were inoculated into *L. perenne* seedlings and the phenotype of infected plants examined eight weeks post-planting. At this time point, WT infected plants typically have 2–6 tillers that are up to 50 cm in length (Fig 7). Plants infected with the deletion mutants had the same interaction phenotype as those infected with WT, with no statistically significant difference in either the number or length of tillers (Fig 7). To analyse the cellular phenotype of the infected plants, pseudostem cross sections were fixed and embedded in resin, and slices stained with Toluidine blue. WT-infected samples typically have one to two hyphae per intercellular space and never colonize the vascular bundle tissue (Fig 8). The mutant-infected plants had the same cellular phenotype as WT plants, with one to two hyphae per intercellular space and no hyphae within the vascular bundles (Fig 8). Given these results, samples were then prepared for TEM, to give better resolution of the cellular phenotype. The cellular structure of tissue infected with *festucae* mutants was indistinguishable from WT (Fig 8). In addition, longitudinal leaf sections were infiltrated with wheat germ agglutinin, a chitin-binding lectin, conjugated to AlexaFluor 488 (WGA-AF488), as well as aniline blue, a β-glucan-binding dye, and analysed by confocal laser scanning microscopy (CLSM). Leaves infected with WT typically had one hypha (red pseudocolor, aniline blue) with regularly spaced septa (blue pseudocolor, WGA-AF488) growing between plant cells (Fig 9). These hyphae occasionally branched and where tips of branches meet, fused to establish a hyphal network throughout the leaf tissue (Fig 9). The morphology and growth pattern of the mutant hyphae were indistinguishable from WT (Fig 9).

**Fig 7.**
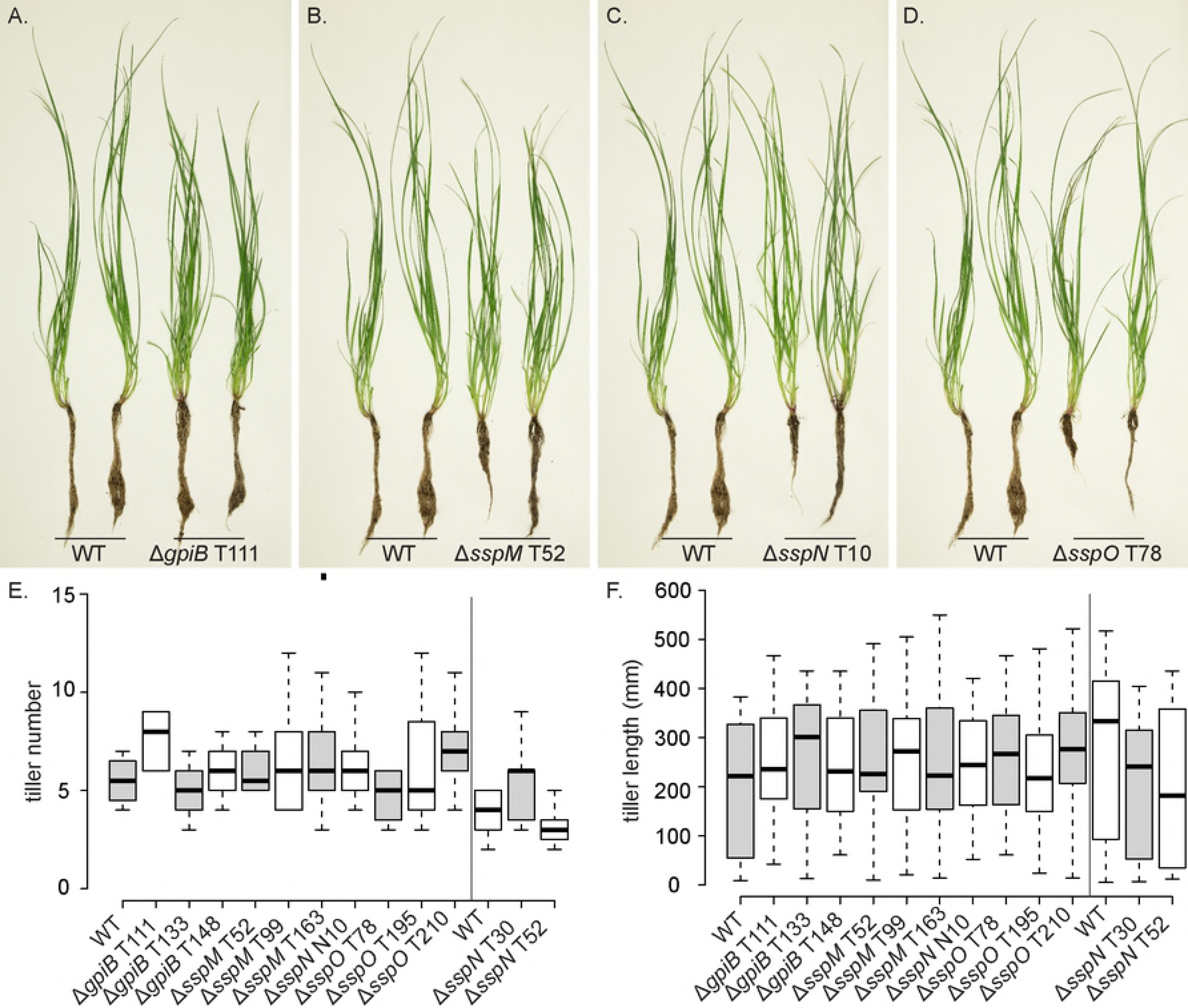
Plant phenotype of *L. perenne* infected with *E. festucae* WT and *gpiB, sspM, sspN* and *sspO* deletion mutant strains. (A-D) Whole plant phenotype of WT and deletion mutant-infected *L. perenne* plants 8 weeks post-planting. (E and F) Box plots representing tiller number (E) and length (F) of ryegrass plants 8 weeks post-planting infected with *E. festucae* WT (n=6) and *ΔgpiB* (n=9/9/5), *ΔsspM* (n=10/15/13), *ΔsspN* (n=13/8/8) and *ΔsspO* (n=7/8/9) mutant strains. One-way ANOVAs were used to test for differences in both plant phenotypes between wildtype and modified strains. In each case the ANOVA was fitted with R and a Bonferroni correction was applied to all p-values to account for multiple testing.

**Fig 8.**
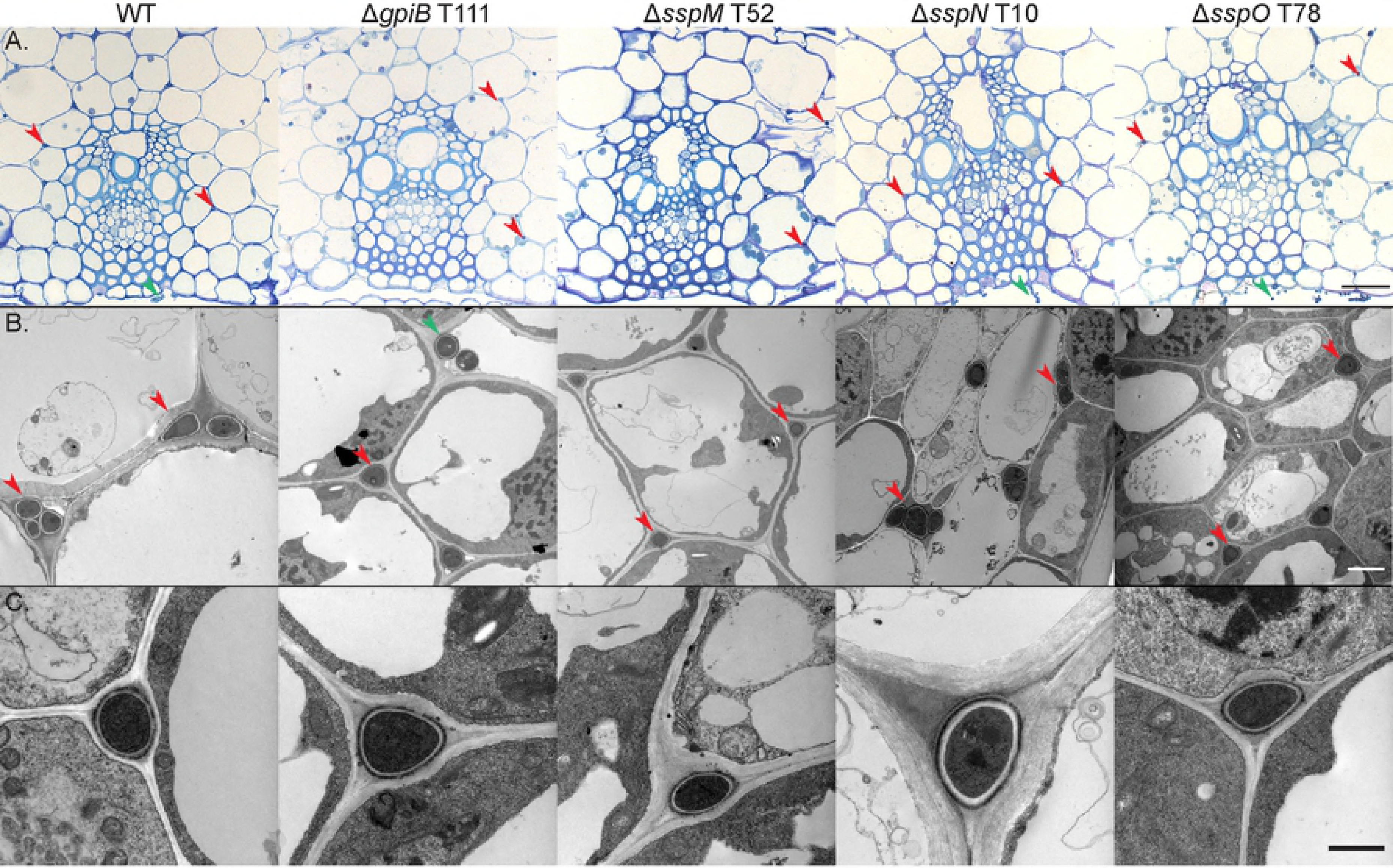
Light microscopy and transmission electron micrographs of the *in planta* cellular phenotype of *E. festucae* WT and *gpiB, sspM, sspN* and *sspO* deletion mutant strains. (A) Light microscope analysis of fixed *L. perenne* pseudostem cross sections stained with Toluidine blue. The focus of the section is on the vascular bundles and surrounding regions; Bar= 10 µm. (B) TEM analysis of hyphal growth in the host apoplast; Bar= 2 µm. (C) Magnified transmission electron micrograph of one single hypha; Bar= 1 µm. Representative images from one of the independent mutants for each gene. Red arrows indicate position of hyphae and green arrows epiphyllous hyphae.

**Fig 9.**
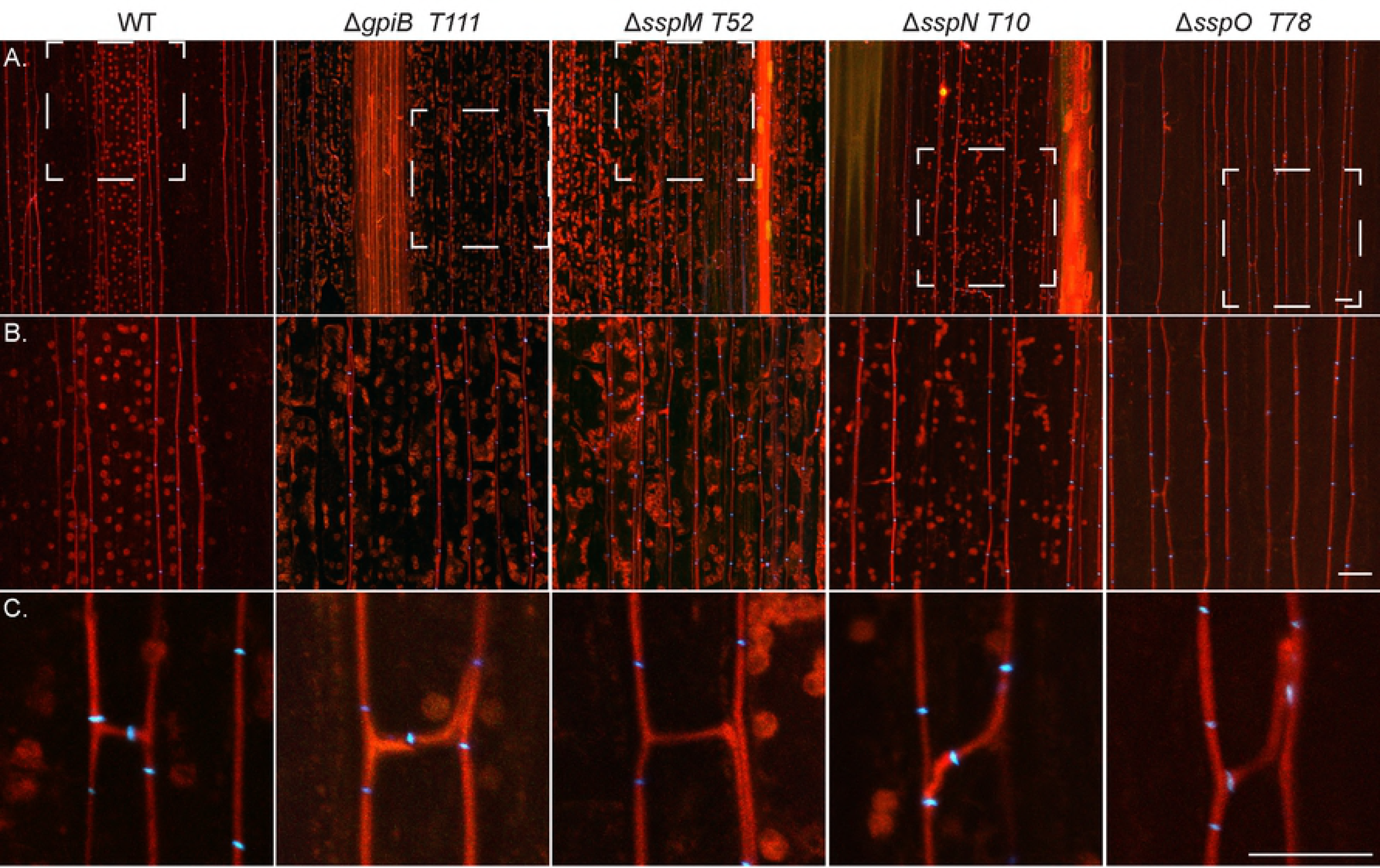
Confocal depth series images of *in planta* cellular phenotype of *E. festucae* WT and *gpiB, sspM, sspN and sspO* deletion mutant strains. Infected *L. perenne* pseudostem samples were stained with aniline blue/WGA-AF488. Hyphae show aniline blue fluorescence of β-glucans (red pseudocolor) and chitin staining of septa with WGA-AF488 (blue pseudocolor). (A) Representative growth of WT and deletion strains *in planta* (*z*= 15 µm); Bar= 20 µm. (B) Magnification of the area boxed with white dashed lines in A, showing representative growth of the strains in more detail (*z*= 15 µm); Bar= 20 µm. (C) Representative hyphal branching and fusion of WT and deletion strains *in planta* (*z*= 5 µm); Bar= 10 µm.

Next we tested whether these mutants could be vertically transmitted into the seeds of *L. perenne*. Plants infected with each of the individual transformants were vernalized and ovaries collected from flowers for CLSM and seed collected from mature inflorescences to test for endophyte in each of these tissues. Endophyte was shown to be present in ovaries of some of the flowers collected from plants infected with Δ*gpiB*, Δ*sspM* and Δ*sspN* mutants but not Δ*sspO* (S3 Fig; S6 Table). Seeds of Δ*gpiB* (2/43), Δ*sspM* (25/90) and Δ*sspN* (14/70) were immunopositive indicating that these three mutants were vertically transmitted (S6 Table). However, none of the seeds infected with Δ*sspO* (0/60) were found to contain endophyte suggesting this mutant was unable to be transferred to the seed.

In summary, deletion of *gpiB, sspM* and *sspN* does not result in any obvious culture or plant interaction phenotype, suggesting that the proteins encoded by these genes are not essential for any of these growth processes. Similarly, *sspO* appears to be dispensable for most of these growth processes but may be important for vertical transmission within the host grass.

### Culture and *in planta* phenotype of overexpression strains

As the deletion of the four genes did not result in any obvious plant interaction phenotype, overexpression (OE) constructs were generated using the translation elongation factor (*tef*) promoter from *Aureobasidium pullulans*, which is commonly used for OE studies [37, 67]. Plasmids encoding the *gpiB, sspM, sspN* and *sspO* OE constructs (pBH29–32) were prepared and transformed into WT protoplasts and the copy number of 10 independent transformants for each construct determined (S4 Fig). A total of three strains per construct, each with a different copy number, were chosen for further analysis. In addition, the relative expression level of the gene in each of the three independent strains was determined and found to generally correlate with the copy number (S4 Fig). No difference in culture growth or morphology was observed between the OE strains and WT. To test the plant symbiotic phenotype, the OE strains were inoculated into *L. perenne* seedlings and infected plants examined 8 weeks post-planting. No difference in whole plant (Fig 10) or cellular phenotype (Fig 11) was observed between the OE strains and WT. In summary, these results demonstrate that overexpression of *gpiB, sspM, sspN* or *sspO* has no impact on the culture or whole plant interaction phenotype.

**Fig 10.**
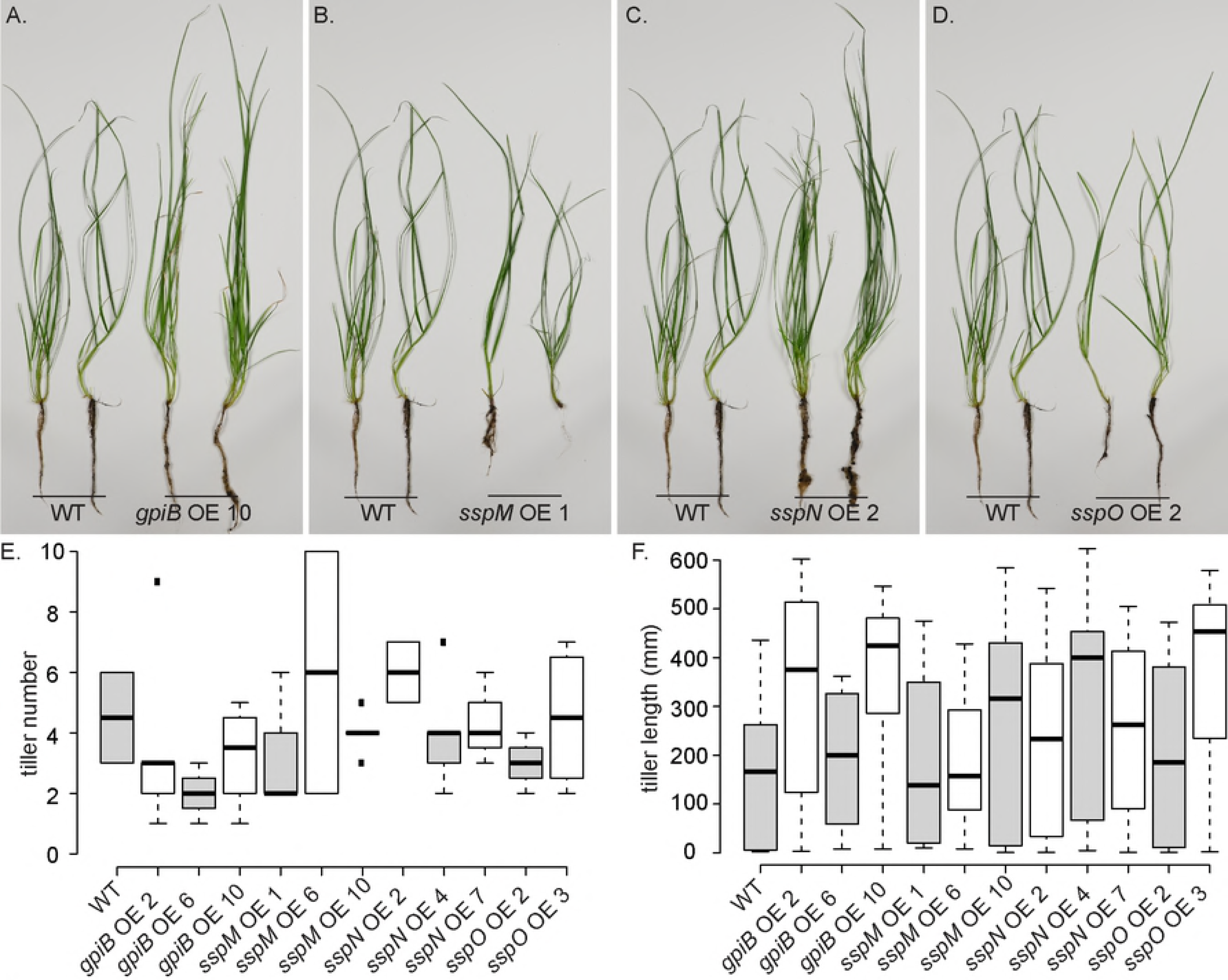
Plant phenotype of *Lolium perenne* infected with *E. festucae* WT and *gpiB, sspM, sspN* and *sspO* OE strains. (A-D) Whole plant phenotype of WT and OE strain-infected *L. perenne* plants 8 weeks post-planting. (E and F) Boxplots representing tiller number (E) and length (F) of ryegrass plants eight weeks post-planting with *E. festucae* WT (n= 2) and *gpiB* (n= 2/6/4), *sspM* (n= 3/2/6), *sspN* (n= 2/7/3) and *sspO* (n= 4/4) OE strains. One-way ANOVAs were used to test for differences in both plant phenotypes between wildtype and modified strains. In each case the ANOVA was fitted with R and a Bonferroni correction was applied to all p-values to account for multiple testing.

**Fig 11.**
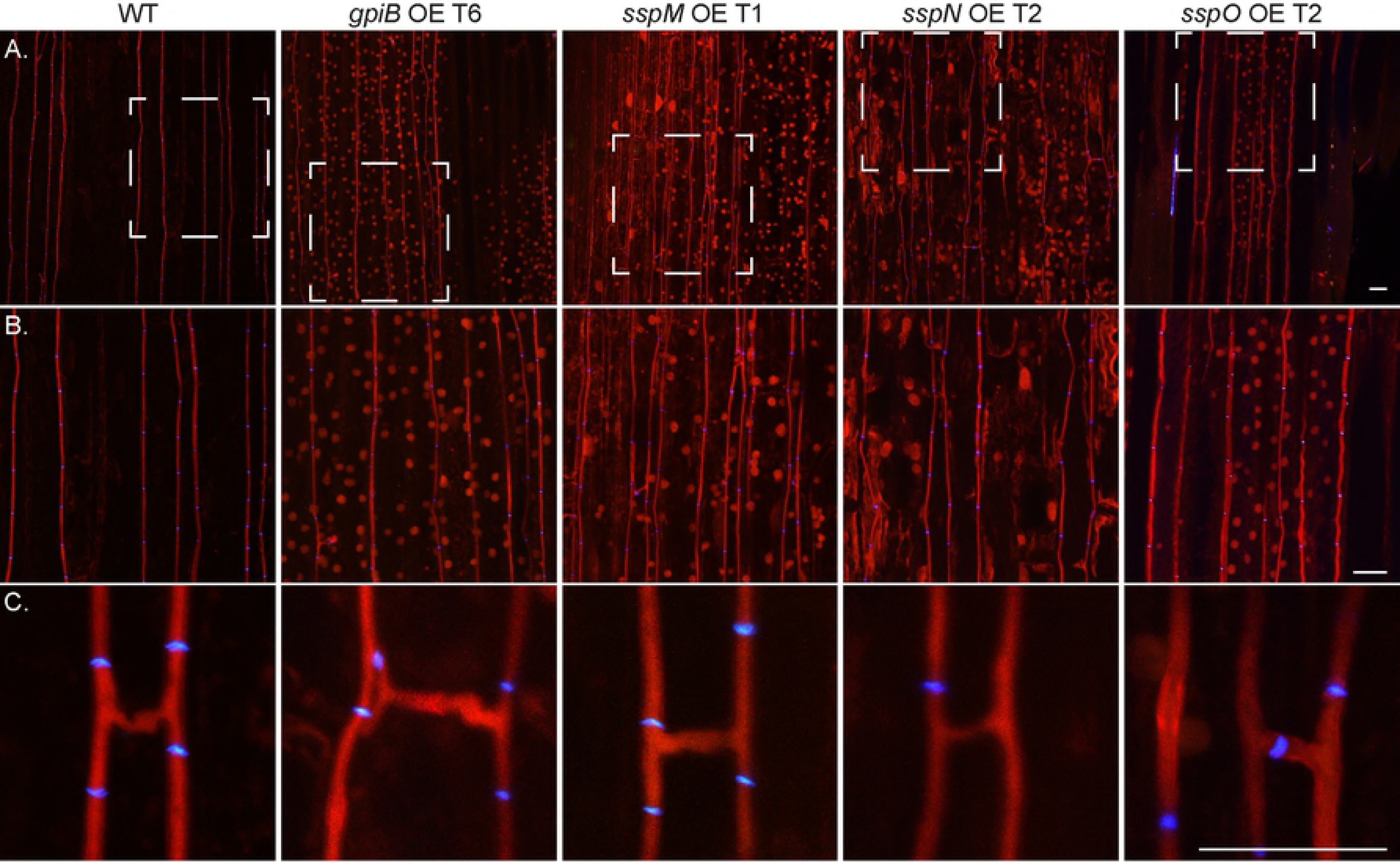
Confocal depth image series of *in planta* cellular phenotype of *E. festucae* WT and *gpiB, sspM, sspN and sspO* OE strains. Infected *L. perenne* pseudostem samples were stained with aniline blue/WGA-AF488. Hyphae show aniline blue fluorescence of β-glucans (red pseudocolor) and chitin staining of septa with WGA-AF488 (blue pseudocolor). (A) Representative growth of WT and OE strains *in planta* (*z*= 15 µm); Bar= 20 µm. (B) Magnification of the area boxed with white dashed lines in A, showing representative growth of the strains in more detail (*z*= 15 nm); Bar= 20 µm. (C) Hyphal branching and fusion of WT and OE strains *in planta* (*z*= 5 nm); Bar= 20 µm.

### SspM, SspN and SspO localize extracellularly

To test whether SspM, SspN and SspO are translocated to the cytosol of the host, we generated C-terminal mCherry fusion constructs of each protein under the control of their native promoters. These constructs also contained three C-terminal tandem repeats of the simian virus large T-antigen nuclear localization signal to target the fusion proteins to the host nucleus, to enhance the signal in the cell if the proteins are translocated to the plant cytosol [60]. The constructs were transformed into protoplasts of their respective deletion strain, together with a vector encoding cytoplasmic eGFP to facilitate identification of hyphae among the plant cell background. Transformants were screened by PCR for presence of the construct and then inoculated into *L. perenne* seedlings. Leaf tissue was harvested from mature infected plants and examined by CSLM for mCherry localization. Although the fluorescence from the mCherry fusion proteins was generally weak *in planta*, a signal of sufficient intensity was detected from SspM-mCherry-NLS and SspO-mCherry-NLS in the apoplast on either side of the hyphae. For SspN-mCherry-NLS, the signal was so weak that a localization profile for this protein in endophytic hyphae was not possible. No evidence was found for localization of any of the three constructs to the plant nucleus despite examining many individual sections and fields from different plants inoculated with other transformants. By contrast, the signal for mCherry in epiphyllous hyphae on the leaf surface was relatively strong and distinctly localized along the cell walls for all three constructs, demonstrating that the proteins were secreted *in planta,* but appear to remain attached to the hyphal cell surface (Fig 12). These observations indicate that SspM, SspN and SspO are secreted into the apoplast and may remain attached to the hyphal cell surface post-secretion.

**Fig 12.**
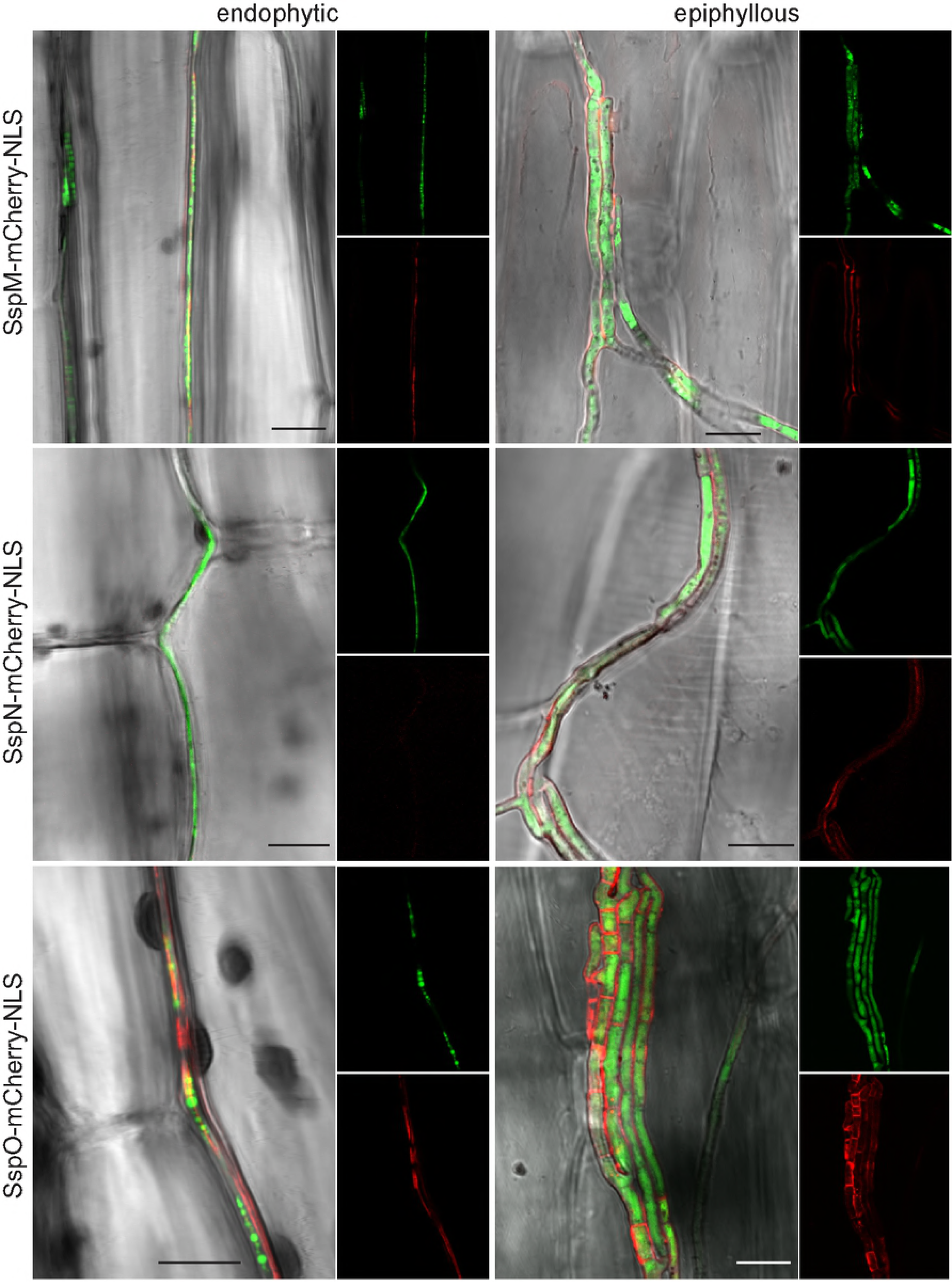
Confocal micrographs of the localization of SspM-mCherry-NLS, SspN-mCherry-NLS and SspO-mCherry-NLS in endophytic and epiphyllous hyphae of *E. festucae* in mature *L. perenne* plants. Besides the mCherry constructs each transformant contained eGFP to facilitate identification of the hyphae *in planta*; Bar= 10 µm.

### SspM, SspN, SspO and GpiB do not elicit a defense response in *N. benthamiana* or *N. tabacum* and do not suppress INF1-triggered immunity in *N. benthamiana*

Several fungal effector proteins have been shown to activate the plant immune system (i.e. to trigger a cell death immune response) in non-host plants upon their recognition as ‘invasion patterns’ by corresponding plant immune receptor proteins [95]. To test whether non-host plants are able to recognize GpiB or the Ssps, as invasion patterns, each was expressed in *N. benthamiana* and *N. tabacum* leaf tissue using an *A. tumefaciens* transient expression assay and screened for an associated cell death response. For this purpose, the cDNA sequence encoding each mature protein (i.e. lacking a native secretion signal) was fused to the nucleotide sequence encoding the *N. tabacum* PR1α secretion signal (for secretion to the plant apoplast) and 3xFLAG (for detection by western blot), and ligated into the pICH86988 expression vector by Golden Gate cloning. The resulting expression vectors were then transformed into *A. tumefaciens* strain GV3101. Strains confirmed to contain the correct construct were infiltrated into leaves of approx. five-week old *N. benthamiana* and *N. tabacum* plants, and the expression of GpiB and the three Ssps verified by western blotting using an antibody to the FLAG-Tag (S5 Fig). Plants were examined for any sign of a cell death response at one-week post-infiltration. As expected, the positive control, INF1 from *Phytophthora infestans* (a PAMP), triggered a strong cell death response in both *N. benthamiana* and *N. tabacum* (Figs 13A and B). As expected, the negative control, pICH86988 empty vector, failed to trigger cell death in these plants (Figs 13A and B). Unlike INF1, however, neither GpiB, nor the three Ssps, triggered cell death in *N. benthamiana* and *N. tabacum* (Figs 13A and B) suggesting that they are not recognized as invasion patterns by these non-host plants.

**Fig 13.**
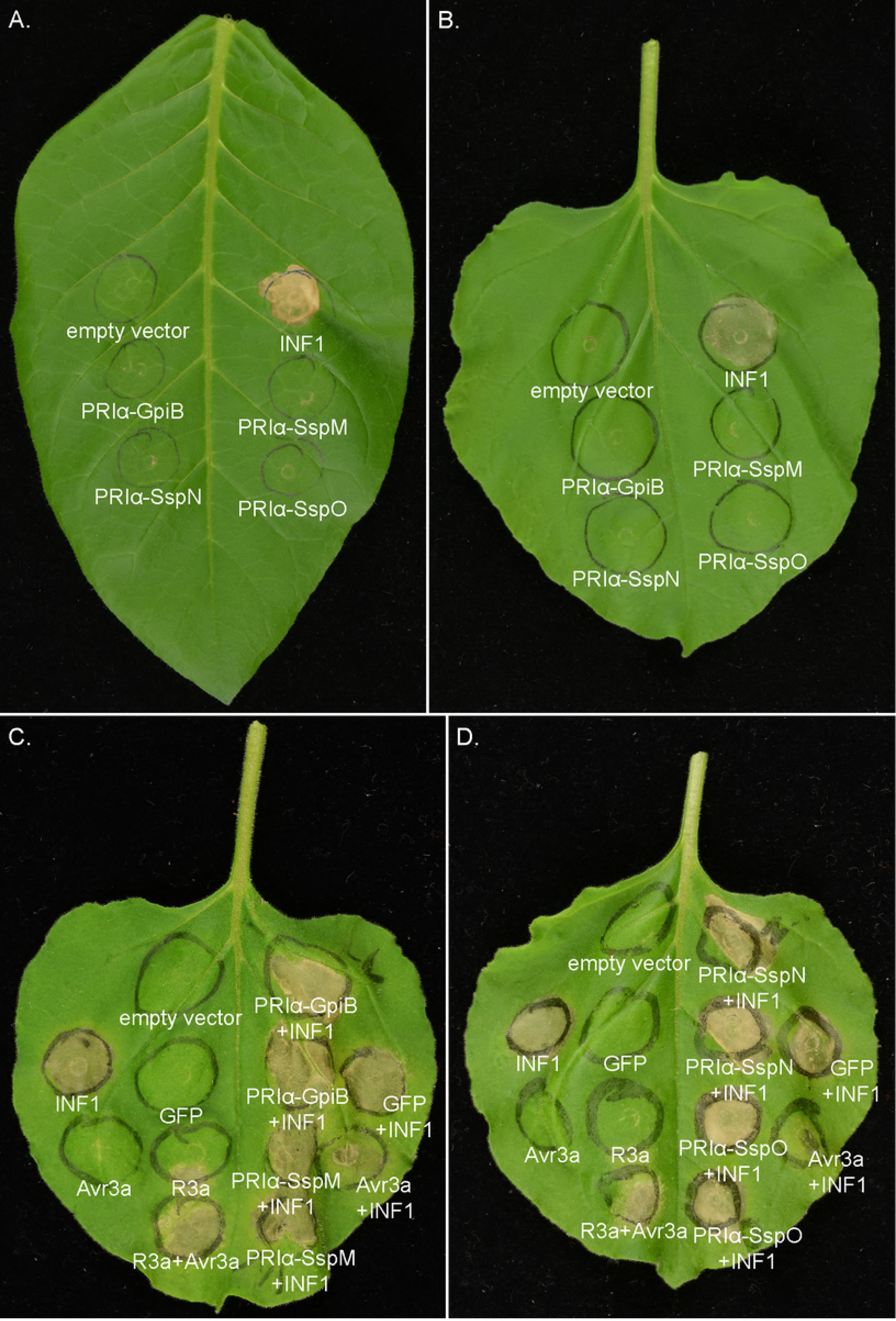
*A. tumefaciens* transient expression assays of GpiB, SspM, SspN and SspO in the non-host plants *N. benthamiana* and *N. tabacum*. (A) *N. benthamiana* and (B) *N. tabacum* leaf phenotypes in response to *A. tumefaciens* transient expression assays involving GpiB, SspM, SspN, SspO or INF1. INF1 triggers a strong cell death response, whereas the pICH86988 empty vector had no effect. (C and D) INF1-triggered cell death suppression assays in *N. benthamiana*. GpiB or any of the three Ssps were unable to suppress INF1-triggered cell death in *N. benthamiana*. eGFP was used as a negative control, INF1/Avr3a as a suppression control, and Avr3a/R3a as cell death positive control. All photos were taken one-week post-infiltration.

Many fungal effector proteins are anticipated to suppress activation of the plant immune response [1]. With this in mind, we next examined whether any of the four proteins are able to suppress the cell death response elicited by the INF1 PAMP in *N. benthamiana*, a model plant system for testing for this response. As before, responses were evaluated one-week post-infection. For this experiment, leaves were first infiltrated with the *A. tumefaciens* strain containing an expression vector for GpiB or one of the three Ssps. After 24 h, the same infiltration zone was infiltrated with the *A. tumefaciens* strain containing the INF1 expression vector. As the outcome of this experimental design is very sensitive to environmental influences, appropriate controls were included for each independent experiment. As expected, infiltration of an *A. tumefaciens* strain containing an expression vector for the *P. infestans* effector Avr3a 24 h after the corresponding potato resistance protein R3 resulted in a strong cell death response (Figs 13C and D). As expected, infiltration of an *A. tumefaciens* strain containing an expression vector for eGFP 24 h before INF1, gave a strong cell death response, demonstrating that eGFP is not able to suppress the cell death response caused by INF1 (Figs 13C and D). When Avr3 was infiltrated 24 h before INF1, the PAMP-triggered cell death response normally elicited by INF1 was strongly suppressed (Figs 13C and D), confirming a previous study showing that Avr3a is able to suppress INF1-triggered cell death in *N. benthamiana* [96]. Unlike the results shown for Avr3a, infiltration of the three Ssps or GpiB, did not result in suppression of INF1-triggered cell death (Figs 13C and D).

Taken together, these experiments show that neither the Ssps, nor GpiB, are recognized by the non-host plants *N. tabacum* and *N. benthamiana*, and that these proteins are unable to suppress the cell death response triggered by INF1.

## Discussion

Effectors of plant-pathogenic fungi function to promote host colonization, typically by suppressing the plant immune system. Current evidence, based on a small number of functionally characterized effectors from root endophytes [16-19], suggests that effectors from plant-mutualistic fungi play a similar role. In this study, we identified a suite of candidate effectors from the mutualistic symbiotic fungus *E. festucae*. We then performed an in-depth functional analysis in an attempt to uncover a role for three candidate effectors, representing some of the most highly expressed candidate effectors from this fungus *in planta*, in the mutualistic symbiotic interaction of *E. festucae* with *L. perenne*.

Making use of the completely annotated genome of *E. festucae* strain Fl1 and a combination of SignalP, EffectorP and protein size restriction we identified 141 genes encoding putative effectors. Eighty-three of these genes are homologous to uncharacterized proteins from other Sordariomycete genera, while a further nine have homologs in other *Epichloë* species. The repertoire of effectors present in the genome thus includes conserved proteins that may support somewhat generalized host-adaptation and a number of apparently lineage-specific genes. Effectors are known to contribute to host specificity in other plant-associated fungal species [97-99], and we predict some of the apparently lineage-specific effectors identified here will contribute to specific host interactions in *Epichloë* strains.

Having a complete and finished *E. festucae* genome sequence enabled us to test for association between structural features of the genome and genes encoding effectors. Our results are remarkable for the lack of associations that are well-documented in other fungal species. The *E. festucae* genome has a “patchwork” structure, in which distinct blocks of highly AT-biased DNA comprised almost entirely of TEs are interleaved with gene-rich regions that have balanced GC-content. This genome structure is similar to that found in the pathogenic fungus *L. maculans,* where effector genes are greatly over-represented within AT-rich blocks (comprising 80% of all genes in these regions). Similarly, it has been reported that effector-encoding genes are located in TE-rich regions of the *M. oryzae* [100], *Blumeria graminis* [101] and *oxysporum* [102] genomes [103]. In contrast, we found no evidence for enrichment of genes encoding candidate effectors in proximity to AT- and transposable element-rich regions. Indeed, only 5.7% of the candidate effector genes can be found within 1 kb of an AT-rich block and none were found within an AT-rich block.

The lack of association between AT-rich blocks in the *E. festucae* genome and effectors is made more surprising by the evidence that these blocks contribute to plant-specific gene expression of other genes [26]. We have recently shown that the patchwork structure of the *E. festucae* genome strongly influences the resultant 3D conformation of the chromosomes, where AT-rich blocks mainly interact with other AT-rich blocks to generate a highly condensed chromatin state compared to the GC and gene-rich blocks. Genes specific to *E. festucae* as well as genes highly expressed during the interaction with the plant host were found to be overrepresented in proximity to the AT-rich blocks [26]. The fact effectors are not associated with the blocks, the rapidly evolving sub-telomeric regions, and do not form distinct clusters as found in other species, reflect the difference in evolutionary pressures experience by symbiotic, rather than pathogenic, fungal species. However, we did find an enrichment of genes encoding secreted proteins in general and SSPs in particular, in proximity to MITEs. MITEs are small, non-autonomous DNA transposable elements abundantly distributed in eukaryotic genomes. Transposable elements are increasingly seen as important for gene regulation, and MITEs have been linked to gene expression traits in a number of species [27, 104]. In *E. festucae*, MITEs have been found in the promoter region of key secondary metabolite gene clusters [24, 105] and are greatly over-represented in the regions upstream of genes preferentially expressed *in planta* [26]. These results, and the demonstration here that genes encoding Ssps are frequently in close proximity to MITEs, suggests that elements either directly or indirectly associated with MITEs may play a key role in regulating the expression of *ssps*.

In the root endophyte *F. oxysporum,* most effector genes are encoded on a dispensable TE-rich pathogenicity chromosome [97, 106]. Recently a MITE from the mimp family was identified in the promoter region of each of the *SIX* effector genes and based on the presence of this mimp in the promoter region of other genes, further secreted proteins could be predicted and subsequently identified in the xylem sap of infected tomato plants [25]. Interestingly, deletion of the mimp element from the promoter region of two of the *SIX* genes did not significantly change the expression of these genes *in planta* or culture, thereby ruling out a direct role of this element in transcriptional regulation [25]. However, a putative transcriptional regulator (repressor) binding element in close proximity to the mimp is important for SIX gene expression. Whether the MITEs identified in *E. festucae* contribute to transcriptional silencing or activation remains to be elucidated.

Expression of effector genes is often induced upon plant infection or at a certain stage of host infection [87, 88]. Consistent with this observation we found a preferential expression of candidate effector-encoding genes *in planta.* Previous work has shown that a major remodeling of the *E. festucae* chromatin state occurs upon the switch from growth in axenic culture to growth *in planta*. In culture the genome is highly heterochromatic, characterized by high levels of H3K9me3 and H3K27me3, whereas *in planta* the genome contains higher levels of transcriptionally active euchromatin characterized by much lower levels of these two repressive epigenetic marks [68]. While RNAseq data for mutants of the responsible methyltransferases was not available, a set of data was available for a mutant of the gene encoding heterochromatin protein 1 (*hepA*). When a comparison was made between the transcriptome of Δ*hepA* vs WT in axenic culture there was a minor but significant increase in expression of the candidate effector genes. Similar results have been obtained in *L. maculans* where silencing of genes encoding the heterochromatin regulators DIM-5, a H3K9 methyl transferase, and heterochromatin protein 1 (HP1) resulted in the expression of many effector genes in axenic culture, which were repressed in WT [89]. As discussed above, effector genes in *L. maculans* are mainly located within the AT isochore component of the genome, which is in a highly heterochromatic state in axenic culture [89]. In contrast genes encoding *E. festucae* effectors are not associated with the AT-rich blocks of the genome and thus were expected to be less affected by heterochromatin silencing.

Previous studies have shown that the deletion of genes coding for NoxA, a NADPH oxidase [34], ProA, a transcription factor [39], and SakA, a stress-responsive MAP kinase [36], in *E. festucae* result in a severe breakdown in the symbiotic interaction with the host leading to severely stunted plants. Analysis of the fungal transcriptome of these three different interactions identified a core set of 182 *E. festucae* genes that were differentially expressed (DE) in the same direction in all three symbiosis mutants [47]. Among this core gene set were 14 genes encoding secreted proteins, corresponding to 10 that were upregulated and four (*gpiB, sspM, sspN* and *sspO*), that were downregulated. The three putative effector genes analysed here were among the top 100 DE genes *in planta* compared to axenic culture.

In order to verify that the predicted secretion signals of the three effector candidates and the GPI-anchored protein were functional, a yeast complementation assay was carried out [52]. *S. cerevisiae* transformants containing each of the four predicted secretion signals fused to SUC2 all grew on raffinose confirming these signals are functional. In addition, western blots were performed with His-tagged versions of each of the four genes under the control of a *tef* promoter to confirm they were secreted and to check for proteolytic processing and any potential post-translational modifications. Both GpiB and SspM were predicted to be *N*-glycosylated and SspM and SspN contain a kexin cleavage site. Proteins of the expected size for GpiB, SspN and SspO were detected in extracts of fungal mycelia. For SspM, two signals were detected which potentially correspond to the pro-protein and a post-translationally modified protein. Some effectors, such as *C. fulvum* Avr4 and Avr9 are further processed in the apoplast by either fungal or plant proteases [20]. To determine whether there is protease modification of the SspM *in planta* and whether this processing is necessary for the activation of SspM will require further experimentation using protein extracts obtained from ryegrass apoplastic fluid. Despite the presence of an LxKR kexin recognition motif in SspN, only a single band was detected corresponding to the size of the unprocessed protein. Sequence alignment with closely related species shows that this motif can only be found in strain Fl1, suggesting this might be the result of a recent mutation. One possible reason for lack of cleavage at this site could be lack of accessibility of this motif within the folded protein. Alternatively, it is possible it is only cleaved *in planta* and not in axenic culture. While the function of SspN is not known it does have a repeat domain structure which is a common feature of several effectors characterized to date including Ecp6 from *C. fulvum* [107], which contains multiple LysM domain containing repeats, Sp7 from the mutualist *Glomus intraradices* [18], *Colletotrichium graminicola* EP1[108] and the recently identified effector Rsp3 from *U. maydis* [6]. However, unlike all these proteins, which have been shown by genetic analysis to be very important for the interaction with the host, deletion of *sspN* had no obvious plant interaction phenotype. Some of these repeat-containing effectors function in the apoplast (Ecp6 and Rsp3), whereas others are transported into the host cytoplasm (Sp7 and EP1). Due to the adoption of an extended module-like structure, some repeat-containing proteins such as EP1 are found to be associated with nucleic acids following transport into the host nucleus whereas others mediate protein-protein interactions [108-110]. In addition, some repeat-containing proteins are located on the surface of the cell where they are integrated into the plasma membrane or attached to the cell wall [109]. Rps3 is located on the *U. maydis* cell wall where it interacts with the mannose-binding and anti-fungal activity of the maize protein AFP1 [6]. Some repeat-containing proteins contain kexin cleavage sites between the repeating units resulting in generation of bioactive peptides. A classic example of this in *Epichloë* is GigA, which is processed to generate a series of cyclic peptides called epichloëcyclins of unknown biological function [111]. In *U. maydis*, Rep1, is processed into smaller peptides that form cell wall associated fibrils mediating attachment of the hyphae to hydrophobic surfaces [112, 113].

Although products of all four genes could be detected in the fungal mycelium, only SspM and SspO, which contain six and eight cysteines respectively, were detected in the culture medium. The lack of detection of GpiB in the culture medium was not surprising as the addition of a GPI-anchor generally results in cell wall attachment. GPI-anchored proteins are involved in multiple processes, including attachment of hyphae to various surfaces, cell wall integrity and virulence [42, 43]. The two protein products observed for SspM in the media were the same size as those observed in the mycelia. The presence of two bands could either indicate that both the modified as well as the unmodified protein fulfill a function in *E. festucae* or that the overexpression of the gene because of multiple integrations resulted in an overproduction compared to the available protease. As the protein however can be detected in the modified version in culture and only one band can be detected in the western blot on SspM expressing *N. benthamiana* protein extracts, a modification by plant proteases to induce activity can probably be excluded. The inability to detect SspN in the culture media could be either due to the instability of the protein resulting from the lack of cysteines that promote stability or due to an attachment of this protein to the cell wall as has been described for multiple repeat-containing proteins as discussed above. A single protein product of the same size as observed in mycelia was detected in the media of cultures containing the SspO construct indicating this protein is not post-translationally modified. The presence of eight cysteines, which have the potential to form four cysteine bridges, is likely to account for the high stability of this protein. Given the observed size for SspM was similar to the expected size, this proteins is probably not *N*-glycosylated. However the product of the His-tagged GpiB was greater than the predicted size, which may be due to addition of the GPI anchor and/or modification by *N*-glycosylation. For a definitive answer, purified proteins could be analyzed by mass spectrometry or subjected to an enzymatic removal of glycosylation followed by a size comparison using SDS-PAGE.

Some pathogens such as *U. maydis* produce apoplastic as well as cytoplasmic effectors [1], whereas other pathogens such as *C. fulvum* seem to only produce apoplastic effectors [72]. In order to determine where the effector candidates of *E. festucae* are targeted during host colonization, translational fusions were generated between each of the candidate genes and mCherry. Given it is difficult to detect effectors in the host cytoplasm, we used the method of adding a nuclear localization signal to target the proteins to the plant nucleus, as this was shown to be very effective for detecting some *M. oryzae* effectors in cells of the host *O. sativa* [60]. Despite examining multiple samples of different aged plants, no signal from any of the candidates was observed in the nuclei of ryegrass leaf cells. Therefore, the effector candidates are likely to reside in the apoplast. Indeed, we were able to show that all three of the effector candidates localized to the cell walls or plasma membrane of the fungal hyphae *in planta*. The clearest signals were obtained by imaging epiphyllous hyphae on the surface of the leaves. While it was more challenging to detect the localization of these proteins inside the leaf, signals of sufficient intensity were detected for both SspM and SspO on the cell walls of endophytic hyphae. However, no signal could be detected for SspN in endophytic hyphae which could be due to the fact that the promoter of *sspN* is weaker than that of *sspM* and *sspO*. Alternatively, SspN may be more unstable in the apoplast than SspM and SspO as was suggested by the western analysis. Alternatively, *sspN* may be just expressed on the leaf surface. This hypothesis could be tested by fusing the promoter of *sspN* to eGFP and analyzing where fluorescence occurs.

To elucidate the role of these proteins in the interaction we generated deletion and overexpression strains and analysed their phenotype in axenic culture as well as in the interaction with perennial ryegrass. In culture, *E. festucae* forms white, soft looking colonies with a diameter of approx. 2 cm after 7 days of growth. Both deletion and overexpression strains were indistinguishable from the WT. Within the plant host, *E. festucae* grows exclusively in the apoplastic space, where the growth is highly regulated and synchronized with the host [114]. In the host meristematic tissue, *E. festucae* grows by tip growth, but as hyphal cells grow into the cell expansion zone of the leaf, the hyphae switch to intercalary growth, a pattern of growth that prevents mechanical shear and enables the hyphae to maintain the same rate of growth as the host [114, 115]. Ryegrass plants infected with either the deletion or overexpression strains had the same whole plant interaction and cellular growth phenotype as WT with one exception. Mutants Δ*gpiB*, Δ*sspM* and Δ*sspN* were seed transmitted but not Δ*sspO*. Additional long-term experiments will be necessary to determine the mechanism for this interesting Δ*sspO* phenotype. Also it would be interesting to check whether these mutants affect the transition from asexual to sexual development but this is difficult to investigate as initiation of the sexual cycle requires vernalisation of host material, formation of stromata and transfer of spermatia to a stroma of the opposite mating type by a *Botanophila* fly [116]. Furthermore, it is possible that each of these effector candidates contributes to the interaction in a minor incremental way making it difficult to detect a phenotype by deletion of single effector gene unless there is a very sensitive phenotype screen [117, 118].

Functional redundancy of effector proteins could also explain the lack of an altered host interaction phenotype. Redundancy is thought to be the result of multiple effectors having the same or overlapping functions. Notably, a study by Weßling and colleagues [119] found effectors from a single pathogen species converging on interactions with a small subset of host proteins in a process named “interspecies convergence” of which many are involved in high level regulatory processes. Interestingly, even effectors from different species of pathogens can target a similar subset of host proteins with a correlation between the degree of convergence and the relevance of the host protein. Therefore homology-independent functional redundancy is maintained as an evolutionary benefit and protects against rapid loss of a single effector due to host recognition [119, 120]. For example, both Avr4 and Ecp6 prevent chitin recognition in the *C. fulvum*-tomato interaction, however, the molecular functions are different and there is no sequence homology between these two effectors [2, 4].

In the presence of a corresponding host immune receptor protein, fungal effectors are known to activate the plant immune system. Often, the main output of this activation is the hypersensitive response (HR), which renders the host resistant and the pathogen avirulent [121]. Notably, a subset of fungal effectors can also trigger an HR-like cell death response when expressed in the non-host plants *N. benthamiana* or *N. tabacum* (i.e. following their recognition by unknown immune receptor proteins) using an *A. tumefaciens* transient expression assay. Indeed, multiple effectors of the wheat pathogen *Z. tritici* have been found to induce an HR in *N. benthamiana* [95]. Using this approach, we tested whether the three candidate effectors of *E. festucae* trigger cell death when produced by these two non-host plants but none triggered a cell death response. We also tested whether the effector candidates are able to suppress a defense response triggered by the *P. infestans* elicitor INF1 using an *A. tumefaciens* transient expression assay. INF1 functions in the apoplastic space where it binds to the ELR (elicitin response) receptor, resulting in activation of the plant immune system [122, 123]. Even though these candidate effectors localize to the apoplast, none were able to suppress INF1-triggered immunity.

In *E. festucae*-infected *L. perenne* plants, approx. one third of *L. perenn*e genes were found to be differentially expressed compared to non-infected plants. These changes were found to be in the key areas of ‘stress response’, ‘primary metabolism’ and ‘secondary metabolism’ [124]. Genes in the category ‘primary metabolism’ and ‘stress response’, including PR genes, were found to be mainly downregulated, and genes involved in secondary metabolism were found to be upregulated [124]. These findings, together with the fact that these candidate effectors are so highly expressed in *E. festucae in planta*, suggests that these proteins are likely to have a key role in the interaction. Furthermore, microscopic studies in infected *L. perenne* plants revealed a lack of chitin, a crucial PAMP, in endophytic hyphae [124, 125]. This indicates that chitin is either masked, potentially by a similar mechanism as has been described for Avr4 of *C. fulvum,* or modified.

In this study, we have identified for the first time, a suite of effector candidates from the grass endophyte *E. festucae*. Genes encoding effector candidates were found to associate with MITEs but not with AT-rich regions or TEs, indicating that they might not be under strong selection pressure. Three effector candidates were functionally analysed but were found to be dispensable for the interaction with *L. perenne* under the growth conditions analysed. Although we could not find any evidence for a role for any of these effectors in the interaction, the list of effector candidates identified provides a good database for selecting other proteins and strategies for functional analysis. Sequencing of additional *Epichloë* species will help identify effector genes conserved across species or unique to one species and thereby provide important insights into the role of these effectors in host specificity in this agriculturally important symbiosis.

## Acknowledgements

This research was supported by grants from the Tertiary Education Commission to the Bio-Protection Research Centre, the Royal Society of New Zealand Marsden Fund (contract MAU1301) and by Massey University. The authors thank Jordan Taylor, Niki Murray and Matthew Savoian (Manawatu Microscopy and Imaging Centre, Massey University) for assistance with microscopy, Arvina Ram and Karolin Warnecke for technical assistance, Christopher Schardl for provision of the *Epichloë* database sequences, Melissa Guo for provision of strains and advice on *Agrobacterium* infiltration assays, and Sophian Kamoun and Barbara Valent for provision of plasmids.

## Author contributions

**Conceptualization**: BH DW CHM CJE BS.

**Data curation**: BH DW.

**Formal analysis**: BH DW CHM.

**Funding acquisition**: BS.

**Investigation**: BS DW.

**Methodology**: BH DW YB CHM CJE BS.

**Project Administration**: BS.

**Resources**: BS.

**Supervision**: CHM CJE BS.

**Writing – original draft**: BH DW CHM BS.

**Writing – review & editing**: CHM CJE BS.

## Supporting information

### S1 File. Methods for computational analysis

**S1 Fig. Amino acid sequence alignments of E. festucae GpiB, SspM, SspN and SspO.** Alignments of GpiB (A), SspM (B), SspN (C) and SspO (D) with homologs from a selection of *Epichloë* species. Red shading: Protease cleavage site; Green shading: omega site; Purple shading: *N*-glycosylation site; Orange shading: cysteine residues. Protein sequences for *E. baconii, E. elymi* and *E. typhina* were obtained from the Kentucky Endophyte database when available or predicted using FGENESH using *Claviceps purpurea* parameters. The deduced amino acid sequence was used for the alignment. Sequences marked with * were manually annotated.

**S2 Fig. Strategy for deletion of the *E. festucae gpiB, sspM, sspN* and *sspO* genes and confirmation by Southern analysis.** (A) Physical map of the *gpiB* WT genomic locus, linear insert of the *gpiB* replacement construct, pBH1, and the recombinant locus showing restriction enzyme sites for *Nde*I. Grey shading indicates regions of recombination. Numbers indicate the PCR primer pairs used for Gibson assembly (BH1/BH5) and deletion mutant screening (BH43/BH44). (B) PCR screening of deletion candidates with the PCR primer pair BH43/BH44, generated expected bands of 3,997 bp in WT, and 4,602 bp in deletion mutants (C) NBT/BCIP-stained Southern blot of digests (approx. 1 µg) from *E. festucae* WT, Δ*gpiB* T27, *ΔgpiB* T111 (PN3113), Δ*gpiB* T133 (PN3114), Δ*gpiB* T148 (PN3115), Δ*gpiB* T160 and Δ*gpiB* T204 strains probed with digoxigenin (DIG)-11-dUTP-labeled linear insert of pBH1 amplified with the primer pair BH1/BH5. Expected bands of 5,317 bp in WT, and 3,971 bp and 1,951 bp in the deletion mutant.

(D) Physical map of the *sspM* WT genomic locus, linear insert of the *sspM* replacement construct, pBH2, and the recombinant locus showing restriction enzyme sites for *EcoR*I. Grey shading indicates regions of recombination. Numbers indicate the PCR primer pairs used for Gibson assembly (BH9/BH12) and deletion mutant screening (BH45/BH46). (E) PCR screening of deletion candidates with the PCR primer pair BH45/BH46, generated expected bands of 3,615 bp in WT, and 4,190 bp in deletion mutants. (F) NBT/BCIP-stained Southern blot of digests (approx. 1 µg) of *E. festucae* WT, Δ*sspM* T180, Δ*sspM* T173, Δ*sspM* T163 (PN3118), Δ*sspM* T135, Δ*sspM* T99 (PN3117) and Δ*sspM* T52 (PN3116) strains probed with digoxigenin (DIG)-11-dUTP-labeled linear insert of pBH2 amplified with the primer pair BH9/BH12. Expected bands of 2,205 bp and 1,569 bp in WT and 4,349 bp in the deletion mutant.

(G) Physical map of the *sspN* WT genomic locus, linear insert of the *sspN* replacement construct, pBH3, and the recombinant locus showing restriction enzyme sites for *Bam*HI. Grey shading indicates regions of recombination. Numbers indicate the PCR primer pairs used for Gibson assembly (BH13/BH16) and deletion mutant screening (BH47/BH48). (H and J) PCR screening of deletion candidates with the PCR primer pair BH47/BH48, generated expected band of 3,403 bp in WT, and 4,250 bp in deletion mutants. (I) NBT/BCIP-stained Southern blot of digests (approx. 1 µg) of *E. festucae* WT, *ΔsspN* T151 and *ΔsspN* T10 (PN3119), strains probed with digoxigenin (DIG)-11-dUTP-labeled linear insert of pBH1 amplified with the primer pair BH13/BH16. (K) NBT/BCIP-stained Southern blot of digests (approx. 1 µg) of *E. festucae* WT, *ΔsspN* T52 (PN3121), *ΔsspN* T15, *ΔsspN* T103, *ΔsspN* T51, *ΔsspN* T30 (PN3120) and *ΔsspN* T3 strains probed with digoxigenin (DIG)-11-dUTP-labeled linear insert of pBH3 amplified with the primer pair BH13/BH16. Expected bands of 1,710 bp and 3,631 bp in WT, and 1,710 bp and 4,477, in the deletion mutant.

(L) Physical map of the *sspO* WT genomic locus, linear insert of the *sspO* replacement construct, pBH4, and the recombinant locus showing restriction enzyme sites for *Pst*I. Grey shading indicates regions of recombination. Numbers indicate the PCR primer pairs used for Gibson assembly (BH5/BH9) and deletion mutant screening (BH49/BH50). (M) PCR screening of deletion candidates with the PCR primer pair BH49/BH50, generated expected band of 3,654 bp in WT, and 4,482 bp in deletion mutants. (N) NBT/BCIP-stained Southern blot of digests (approx. 1 µg) of *E. festucae* WT, *ΔsspO* T210 (PN3124), *ΔsspO* T195 (PN3123), *Δ*sspO T130, *ΔsspO* T102, *ΔsspO* T78 (PN3122) and *ΔsspO* T60 strains probed with digoxigenin (DIG)-11-dUTP-labeled linear insert of pBH4 amplified with the primer pair BH5/BH9. Expected bands of 1,859 bp and 1,678 bp in WT and 4,635 bp in the deletion mutant.

**S3 Fig. Confocal laser scanning microscopy of aniline blue and WGA-AF488 stained ovaries.** The fungal endophytic cell wall was stained with aniline blue (orange pseudo colour) while fungal septa were stained with WGA-AF 488 (blue pseudo colour). A. Ovary of mutant strain Δ*sspM* T52, B. Ovary of mutant strain Δ*sspM* T163. *E. festucae* hyphae are marked by asterisk. Scale bar: 50 µm.

**S4 Fig. qPCR and RT-qPCR results of the *gpiB, sspM, sspN* and *sspO* OE strains.** (A) Copy number determined by qPCR is expressed relative to the WT copy number. Genes encoding *hepA* (single copy, light grey) and *pacC* (single copy, dark grey) were used as reference genes. (B) Expression level determined by RT-qPCR is expressed relative to the WT gene expression. The 40S ribosomal S22 gene was used as reference gene. Primers used for the analyses are given in S2 Table.

**S5 Fig. Verification of the heterologous production of GpiB, SspM, SspN and SspO in *N. benthamiana* via western blotting.** Total protein of the infiltrated leaf region was extracted and separated by electrophoresis on a 10% SDS gel. The gel was transferred to a membrane and probed with an anti-FLAG antibody. eGFP expressed in *N. benthamiana* served as positive control. Expected sizes: approx. 9.8 kDa for GpiB, 14 kDa for SspN, 10 kDa for SspO and for SspM 14.8 and 18.8 kDa.

S1 Table. Biosample references

S2 Table. Biological material.

S3 Table. Primers used in this study.

S4 Table. Summary of the presented information about *E. festucae* effector candidates.

S5 Table. Distribution of *gpiB* and the *ssps* within *Epichloë* and related species.

S6 Table. Vertical transmission of ssp mutants in *L. perenne*.

## References

1. Lo Presti L, Lanver D, Schweizer G, Tanaka S, Liang L, Tollot M, et al. Fungal effectors and plant susceptibility. Annu Rev Plant Biol. 2015;66:513–45. doi: 10.1146/annurev-arplant-043014-114623. PubMed PMID: 25923844.

2. van den Burg HA, Harrison SJ, Joosten MH, Vervoort J, de Wit PJ. *Cladosporium fulvum* Avr4 protects fungal cell walls against hydrolysis by plant chitinases accumulating during infection. Mol Plant-Microbe Interact. 2006;19(12):1420–30. PubMed PMID: 17153926.

3. van Esse HP, Van’t Klooster JW, Bolton MD, Yadeta KA, van Baarlen P, Boeren S, et al. The *Cladosporium fulvum* virulence protein Avr2 inhibits host proteases required for basal defense. Plant Cell. 2008;20(7):1948–63. Epub 2008/07/29. doi: 10.1105/tpc.108.059394. PubMed PMID: 18660430; PubMed Central PMCID: PMCPMC2518240.

4. de Jonge R, van Esse HP, Kombrink A, Shinya T, Desaki Y, Bours R, et al. Conserved fungal LysM effector Ecp6 prevents chitin-triggered immunity in plants. Science. 2010;329(5994):953–35. Epub 2010/08/21. doi: 329/5994/953 [pii] 10.1126/science.1190859. PubMed PMID: 20724636.

5. Doehlemann G, Reissmann S, Assmann D, Fleckenstein M, Kahmann R. Two linked genes encoding a secreted effector and a membrane protein are essential for *Ustilago maydis*-induced tumour formation. Mol Microbiol. 2011;81(3):751–66. doi: 10.1111/j.1365-2958.2011.07728.x. PubMed PMID: 21692877.

6. Ma LS, Wang L, Trippel C, Mendoza-Mendoza A, Ullmann S, Moretti M, et al. The *Ustilago maydis* repetitive effector Rsp3 blocks the antifungal activity of mannose-binding maize proteins. Nature communications. 2018;9(1):1711. doi: 10.1038/s41467-018-04149-0. PubMed PMID: 29703884; PubMed Central PMCID: PMC5923269.

7. Hurlburt NK, Chen L-H, Stergiopoulos I, Fisher AJ. Structure of the *Cladosporium fulvum* Avr4 effector in complex with (GlcNAc)6 reveals the ligand-binding mechanism and uncouples its intrinsic function from recognition by the Cf-4 resistance protein. PLoS Pathog. 2018;14(8):e1007263. Epub 2018/08/28. doi: 10.1371/journal.ppat.1007263. PubMed PMID: 30148881; PubMed Central PMCID: PMCPMC6128652.

8. Sánchez-Vallet A, Saleem-Batcha R, Kombrink A, Hansen G, Valkenburg D-J, Thomma BPHJ, et al. Fungal effector Ecp6 outcompetes host immune receptor for chitin binding through intrachain LysM dimerization. eLife. 2013;2:e00790. Epub 2013/07/11. doi: 10.7554/eLife.00790. PubMed PMID: 23840930; PubMed Central PMCID: PMCPMC3700227.

9. Mueller AN, Ziemann S, Treitschke S, Assmann D, Doehlemann G. Compatibility in the *Ustilago maydis*-maize interaction requires inhibition of host cysteine proteases by the fungal effector Pit2. PLoS Pathog. 2013;9(2):e1003177. doi: 10.1371/journal.ppat.1003177. PubMed PMID: 23459172; PubMed Central PMCID: PMC3573112.

10. Rooney HCE, Van’t Klooster JW, van der Hoorn RAL, Joosten MHAJ, Jones JDG, de Wit PJGM. Cladosporium Avr2 inhibits tomato Rcr3 protease required for *Cf-2*-dependent disease resistance. Science. 2005;308(5729):1783–6. Epub 2005/04/23. doi: 10.1126/science.1111404. PubMed PMID: 15845874.

11. Shabab M, Shindo T, Gu C, Kaschani F, Pansuriya T, Chintha R, et al. Fungal effector protein AVR2 targets diversifying defense-related cys proteases of tomato. Plant Cell. 2008;20(4):1169–83. Epub 2008/05/03. doi: 10.1105/tpc.107.056325. PubMed PMID: 18451324; PubMed Central PMCID: PMCPMC2390736.

12. Djamei A, Schipper K, Rabe F, Ghosh A, Vincon V, Kahnt J, et al. Metabolic priming by a secreted fungal effector. Nature. 2011;478(7369):395–8. Epub 2011/10/07. doi: nature10454 [pii] 10.1038/nature10454. PubMed PMID: 21976020.

13. O’Connell RJ, Panstruga R. Tête à tête inside a plant cell: establishing compatibility between plants and biotrophic fungi and oomycetes. New Phytol. 2006;171(4):699–718. Epub 2006/08/22. doi: 10.1111/j.1469-8137.2006.01829.x. PubMed PMID: 16918543.

14. Zamioudis C, Pieterse CMJ. Modulation of host immunity by beneficial microbes. Mol Plant-Microbe Interact. 2012;25(2):139–50. Epub 2011/10/15. doi: 10.1094/MPMI-06-11-0179. PubMed PMID: 21995763.

15. Pauwels L, Goossens A. The JAZ proteins: a crucial interface in the jasmonate signaling cascade. Plant Cell. 2011;23(9):3089–100. Epub 2011/10/04. doi: 10.1105/tpc.111.089300. PubMed PMID: 21963667; PubMed Central PMCID: PMCPMC3203442.

16. Plett JM, Kemppainen M, Kale SD, Kohler A, Legue V, Brun A, et al. A secreted effector protein of *Laccaria bicolor* is required for symbiosis development. Curr Biol. 2011;21(14):1197–203. doi: 10.1016/j.cub.2011.05.033. PubMed PMID: 21757352.

17. Plett JM, Daguerre Y, Wittulsky S, Vayssieres A, Deveau A, Melton SJ, et al. Effector MiSSP7 of the mutualistic fungus *Laccaria bicolor* stabilizes the Populus JAZ6 protein and represses jasmonic acid (JA) responsive genes. Proc Natl Acad Sci USA. 2014;111(22):8299–304. doi: 10.1073/pnas.1322671111. PubMed PMID: 24847068; PubMed Central PMCID: PMC4050555.

18. Kloppholz S, Kuhn H, Requena N. A secreted fungal effector of *Glomus intraradices* promotes symbiotic biotrophy. Curr Biol. 2011;21(14):1204–9. doi: 10.1016/j.cub.2011.06.044. PubMed PMID: 21757354.

19. Wawra S, Fesel P, Widmer H, Timm M, Seibel J, Leson L, et al. The fungal-specific beta-glucan-binding lectin FGB1 alters cell-wall composition and suppresses glucan-triggered immunity in plants. Nature communications. 2016;7:13188. doi: 10.1038/ncomms13188. PubMed PMID: 27786272.

20. Stergiopoulos I, de Wit PJGM. Fungal effector proteins. Annu Rev Phytopathol. 2009;47:233–63. Epub 2009/04/30. doi: 10.1146/annurev.phyto.112408.132637. PubMed PMID: 19400631.

21. Rouxel T, Grandaubert J, Hane JK, Hoede C, van de Wouw AP, Couloux A, et al. Effector diversification within compartments of the *Leptosphaeria maculans genome affected by Repeat-Induced Point* mutations. Nature communications. 2011;2:202. doi: 10.1038/ncomms1189. PubMed PMID: 21326234; PubMed Central PMCID: PMC3105345.

22. Raffaele S, Kamoun S. Genome evolution in filamentous plant pathogens: why bigger can be better. Nat Rev Microbiol. 2012;10(6):417–30. Epub 2012/05/09. doi: 10.1038/nrmicro2790. PubMed PMID: 22565130.

23. Feschotte C, Mouches C. Evidence that a family of miniature inverted-repeat transposable elements (MITEs) from the *Arabidopsis thaliana* genome has arisen from a pogo-like DNA transposon. Mol Biol Evol. 2000;17(5):730–7. Epub 2000/04/26. doi: 10.1093/oxfordjournals.molbev.a026351. PubMed PMID: 10779533.

24. Fleetwood DJ, Khan AK, Johnson RD, Young CA, Mittal S, Wrenn RE, et al. Abundant degenerate miniature inverted-repeat transposable elements in genomes of epichloid fungal endophytes of grasses. Genome Biol Evol. 2011;3:1253–64. Epub 2011/09/29. doi: evr098 [pii] 10.1093/gbe/evr098. PubMed PMID: 21948396; PubMed Central PMCID: PMC3227409.

25. Schmidt SM, Houterman PM, Schreiver I, Ma L, Amyotte S, Chellappan B, et al. MITEs in the promoters of effector genes allow prediction of novel virulence genes in *Fusarium oxysporum*. BMC Genomics. 2013;14:119. doi: 10.1186/1471-2164-14-119. PubMed PMID: 23432788; PubMed Central PMCID: PMC3599309.

26. Winter DJ, Ganley ARD, Young CA, Liachko I, Schardl CL, Dupont PY, et al. Repeat elements organise 3D genome structure and mediate transcription in the filamentous fungus *Epichloë festucae*. PLoS Genet. 2018;14(10):e1007467. Epub 2018/10/26. doi: 10.1371/journal.pgen.1007467. PubMed PMID: 30356280.

27. Fattash I, Rooke R, Wong A, Hui C, Luu T, Bhardwaj P, et al. Miniature inverted-repeat transposable elements: discovery, distribution, and activity. Genome. 2013;56(9):475–86. Epub 2013/10/31. doi: 10.1139/gen-2012-0174. PubMed PMID: 24168668.

28. Schardl CL, Scott B, Florea S, Zhang D. Epichloe endophytes: clavicipitaceous symbionts of grasses. In: Deising HB, editor. The Mycota Volume V - Plant Relationships. Berlin: Springer-Verlag; 2009. p. 275–306.

29. Leuchtmann A, Schardl CL, Siegel MR. Sexual compatibility and taxonomy of a new species of *Epichloë* symbiotic with fine fescue grasses. Mycologia. 1994;86:802–12.

30. Schardl CL, Leuchtmann A, Tsai H-F, Collett MA, Watt DM, Scott DB. Origin of a fungal symbiont of perennial ryegrass by interspecific hybridization of a mutualist with the ryegrass choke pathogen, *Epichloë typhina*. Genetics. 1994;136:1307–17.

31. Chung K-R, Schardl CL. Sexual cycle and horizontal transmission of the grass symbiont, *Epichloë typina*. Mycol Res. 1997;101(3):295–301.

32. Scott B, Schardl C. Fungal symbionts of grasses: evolutionary insights and agricultural potential. Trends Microbiol. 1993;1(5):196–200.

33. Chung K-R, Hollin W, Siegel MR, Schardl CL. Genetics of host specifictiy in *Epichloë typhina*. Phytopathology. 1997;87(6):599–605.

34. Tanaka A, Christensen MJ, Takemoto D, Park P, Scott B. Reactive oxygen species play a role in regulating a fungus-perennial ryegrass mutualistic association. Plant Cell. 2006;18:1052–66.

35. Takemoto D, Kamakura S, Saikia S, Becker Y, Wrenn R, Tanaka A, et al. Polarity proteins Bem1 and Cdc24 are components of the filamentous fungal NADPH oxidase complex. Proc Natl Acad Sci USA. 2011;108(7):2861–6.

36. Eaton CJ, Cox MP, Ambrose B, Becker M, Hesse U, Schardl CL, et al. Disruption of signaling in a fungal-grass symbiosis leads to pathogenesis. Plant Physiol. 2010;153(4):1780–94. Epub 2010/06/04. doi: pp.110.158451 [pii] 10.1104/pp.110.158451. PubMed PMID: 20519633.

37. Takemoto D, Tanaka A, Scott B. A p67(Phox)-like regulator is recruited to control hyphal branching in a fungal-grass mutualistic symbiosis. Plant Cell. 2006;18(10):2807–21. PubMed PMID: 17041146.

38. Tanaka A, Takemoto D, Hyon GS, Park P, Scott B. NoxA activation by the small GTPase RacA is required to maintain a mutualistic symbiotic association between *Epichloë festucae* and perennial ryegrass. Mol Microbiol. 2008;68(5):1165–78. PubMed PMID: 18399936.

39. Tanaka A, Cartwright GM, Saikia S, Kayano Y, Takemoto D, Kato M, et al. ProA, a transcriptional regulator of fungal fruiting body development, regulates leaf hyphal network development in the Epichloë festucae-*Lolium perenne* symbiosis. Mol Microbiol. 2013;90(3):551–68. doi: doi: 10.1111/mmi.12385.

40. Becker Y, Eaton CJ, Brasell E, May KJ, Becker M, Hassing B, et al. The fungal cell wall integrity MAPK cascade is crucial for hyphal network formation and maintenance of restrictive growth of *Epichloë festucae* in symbiosis with *Lolium perenne*. Mol Plant-Microbe Interact. 2015;28:69–85. doi: doi:10.1094/MPMI-06-14-0183-R.

41. Sperschneider J, Gardiner DM, Dodds PN, Tini F, Covarelli L, Singh KB, et al. EffectorP: predicting fungal effector proteins from secretomes using machine learning. New Phytol. 2016;210(2):743–61. doi: 10.1111/nph.13794. PubMed PMID: 26680733.

42. Rittenour WR, Harris SD. Glycosylphosphatidylinositol-anchored proteins in *Fusarium graminearum*: inventory, variability, and virulence. PLoS One. 2013;8(11):e81603. doi: 10.1371/journal.pone.0081603. PubMed PMID: 24312325; PubMed Central PMCID: PMC3843709.

43. Zhu W, Wei W, Wu Y, Zhou Y, Peng F, Zhang S, et al. BcCFEM1, a CFEM domain-containing protein with putative GPI-anchored site, is involved in pathogenicity, conidial production, and stress tolerance in *Botrytis cinerea*. Frontiers in microbiology. 2017;8:1807. Epub 2017/10/06. doi: 10.3389/fmicb.2017.01807. PubMed PMID: 28979251; PubMed Central PMCID: PMC5611420.

44. Petersen TN, Brunak S, von Heijne G, Nielsen H. SignalP 4.0: discriminating signal peptides from transmembrane regions. Nat Methods. 2011;8(10):785–6. Epub 2011/10/01. doi: nmeth.1701 [pii] 10.1038/nmeth.1701. PubMed PMID: 21959131.

45. R development core team. R: A language and environment for statistical computing. R Foundation for Statistical Computing, Vienna, Austria. 2013;URL https://www.R-project.org/.

46. Quinlan AR. BEDTools: The swiss-army tool for genome feature analysis. Curr Protoc Bioinformatics. 2014;47:11.2. 1–34. Epub 2014/09/10. doi: 10.1002/0471250953.bi1112s47. PubMed PMID: 25199790; PubMed Central PMCID: PMCPMC4213956.

47. Eaton CJ, Dupont P-Y, Solomon PS, Clayton W, Scott B, Cox MP. A core gene set describes the molecular basis of mutualism and antagonism in *Epichloë* species. Mol Plant-Microbe Interact. 2015;28:218–31. doi: doi.org/10.1094/MPMI-09-14-0293-FI.

48. Lachner M, O’Carroll D, Rea S, Mechtler K, Jenuwein T. Methylation of histone H3 lysine 9 creates a binding site for HP1 proteins. Nature. 2001;410(6824):116–20. doi: 10.1038/35065132. PubMed PMID: 11242053.

49. Maison C, Almouzni G. HP1 and the dynamics of heterochromatin maintenance. Nat Rev Mol Cell Biol. 2004;5(4):296–304. doi: 10.1038/nrm1355. PubMed PMID: 15071554.

50. Miller JH. Experiments in Molecular Genetics. New York: Cold Spring Harbor Laboratory Press; 1972.

51. Sambrook J, Fritsch EF, Maniatis T. Molecular cloning: a laboratory manual. Cold Spring Harbor, New York: Cold Spring Harbor Laboratory Press; 1989.

52. Jacobs KA, Collins-Racie LA, Colbert M, Duckett M, Golden-Fleet M, Kelleher K, et al. A genetic selection for isolating cDNAs encoding secreted proteins. Gene. 1997;198(1-2):289–96. PubMed PMID: 9370294.

53. Ausubel FM, Brent R, Kingston RE, Moore DD, Seidman JG, Smith JA, et al. Current Protocols in Molecular Biology. New York.: John Wiley & Sons; 1993.

54. Moon CD, Tapper BA, Scott B. Identification of *Epichloë endophytes in planta* by a microsatellite-based PCR fingerprinting assay with automated analysis. Appl Environ Microbiol. 1999;65:1268–79.

55. Moon CD, Scott B, Schardl CL, Christensen MJ. The evolutionary origins of *Epichloë* endophytes from annual ryegrasses. Mycologia. 2000;92:1103–18.

56. Latch GCM, Christensen MJ. Artificial infection of grasses with endophytes. Ann Appl Biol. 1985;107(1):17–24.

57. Byrd AD, Schardl CL, Songlin PJ, Mogen KL, Siegel MR. The β-tubulin gene of *Epichloë typhina* from perennial ryegrass (*Lolium perenne*). Curr Genet. 1990;18(4):347–54.

58. Gibson DG, Young L, Chuang RY, Venter JC, Hutchison CA, III, Smith HO. Enzymatic assembly of DNA molecules up to several hundred kilobases. Nat Methods. 2009;6(5):343–5. Epub 2009/04/14. doi: nmeth.1318 [pii] 10.1038/nmeth.1318. PubMed PMID: 19363495.

59. Eisenhaber B, Schneider G, Wildpaner M, Eisenhaber F. A sensitive predictor for potential GPI lipid modification sites in fungal protein sequences and its application to genome-wide studies for *Aspergillus nidulans, Candida albicans, Neurospora crassa, Saccharomyces cerevisiae and Schizosaccharomyces pombe*. J Mol Biol. 2004;337(2):243–53. doi: 10.1016/j.jmb.2004.01.025. PubMed PMID: 15003443.

60. Khang CH, Berruyer R, Giraldo MC, Kankanala P, Park SY, Czymmek K, et al. Translocation of *Magnaporthe oryzae* effectors into rice cells and their subsequent cell-to-cell movement. Plant Cell. 2010;22(4):1388–403. Epub 2010/05/04. doi: tpc.109.069666 [pii] 10.1105/tpc.109.069666. PubMed PMID: 20435900; PubMed Central PMCID: PMC2879738.

61. Winston F, Dollard C, Ricupero-Hovasse SL. Construction of a set of convenient *Saccharomyces cerevisiae* strains that are isogenic to S288C. Yeast. 1995;11(1):53–5. Epub 1995/01/01. doi: 10.1002/yea.320110107. PubMed PMID: 7762301.

62. Gietz RD, Woods RA. Transformation of yeast by lithium acetate/single-stranded carrier DNA/polyethylene glycol method. Meth Enzymol. 2002;350:87–96. Epub 2002/06/21. PubMed PMID: 12073338.

63. Koncz C, Martini N, Szabados L, Hrouda M, Bachmair A, Schell J. Specialized vectors for gene tagging and expression studies. In: Gelvin SB, Schilperoort RA, editors. Plant Molecular Biology Manual. Dordrecht: Kluwer Academic; 1994. p. 1–22.

64. Weigel D, Glazebrook J. Transformation of Agrobacterium using the freeze-thaw method. CSH Protoc. 2006;pdb.prot4666.

65. Young CA, Bryant MK, Christensen MJ, Tapper BA, Bryan GT, Scott B. Molecular cloning and genetic analysis of a symbiosis-expressed gene cluster for lolitrem biosynthesis from a mutualistic endophyte of perennial ryegrass. Mol Gen Genomics. 2005;274:13–29.

66. Itoh Y, Johnson R, Scott B. Integrative transformation of the mycotoxin-producing fungus, *Penicillium paxilli*. Curr Genet. 1994;25:508–13.

67. Lukito Y, Chujo T, Scott B. Molecular and cellular analysis of the pH response transcription factor PacC in the fungal symbiont *Epichloë festucae*. Fungal Genet Biol. 2015;85:25–37. doi: 10.1016/j.fgb.2015.10.008. PubMed PMID: 26529380.

68. Chujo T, Scott B. Histone H3K9 and H3K27 methylation regulates fungal alkaloid biosynthesis in a fungal endophyte-plant symbiosis. Mol Microbiol. 2014;92(2):413–34. Epub 2014/02/28. doi: 10.1111/mmi.12567. PubMed PMID: 24571357.

69. Becker Y, Green K, Scott B, Becker M. Artificial inoculation of *Epichloë festucae* into *Lolium perenne*, and visualization of endophytic and epiphyllous fungal growth. Bio-protocol 2018;8:e2990.

70. Florea S, Schardl CL, Hollin W. Detection and isolation of *Epichloë* species, fungal endophytes of grasses. Current protocols in microbiology. 2015;38:19A.1.–A.1.24. Epub 2015/08/04. doi: 10.1002/9780471729259.mc19a01s38. PubMed PMID: 26237108.

71. Spiers AG, Hopcroft DH. Black canker and leaf spot of Salix in New Zealand caused by *Glomerella miyabeana (Colletotrichum gloeosporioides*). Eur J Forest Pathol. 1993;23:92–102.

72. Mesarich CH, Ökmen B, Rovenich H, Griffiths SA, Wang C, Karimi Jashni M, et al. Specific hypersensitive response-associated recognition of new apoplastic effectors from *Cladosporium fulvum* in wild tomato. Mol Plant-Microbe Interact. 2018;31(1):145–62. doi: 10.1094/MPMI-05-17-0114-FI. PubMed PMID: 29144204.

73. Joosten MHAJ, Vogelsang R, Cozijnsen TJ, Verberne MC, de Wit PJGM. The biotrophic fungus *Cladosporium fulvum* circumvents *Cf-4*-mediated resistance by producing unstable AVR4 elicitors. Plant Cell. 1997;9(3):367–79.

74. Luderer R, De Kock MJD, Dees RHL, De Wit PJGM, Joosten MHAJ. Functional analysis of cysteine residues of ECP elicitor proteins of the fungal tomato pathogen Cladosporium fulvum. Mol Plant Pathol. 2002;3(2):91–5. Epub 2002/03/01. doi: 10.1046/j.1464-6722.2001.00095.x. PubMed PMID: 20569313.

75. Kämper J, Kahmann R, Bolker M, Ma LJ, Brefort T, Saville BJ, et al. Insights from the genome of the biotrophic fungal plant pathogen *Ustilago maydis*. Nature. 2006;444(7115):97–101. doi: 10.1038/nature05248. PubMed PMID: 17080091.

76. Brefort T, Tanaka S, Neidig N, Doehlemann G, Vincon V, Kahmann R. Characterization of the largest effector gene cluster of *Ustilago maydis*. PLoS Pathog. 2014;10(7):e1003866. doi: 10.1371/journal.ppat.1003866. PubMed PMID: 24992561; PubMed Central PMCID: PMC4081774.

77. Cambareri EB, Jensen BC, Schabtach E, Selker EU. Repeat-induced G-C to A-T mutations in Neurospora. Science. 1989;244(4912):1571–5. PubMed PMID: 2544994.

78. Gladyshev E. Repeat-induced point mutation and other genome defense mechanisms in fungi. Microbiol Spectrum. 2017;5(4):FUNK-0042-2017. doi: 10.1128/microbiolspec.FUNK-0042-2017. PubMed PMID: 28721856; PubMed Central PMCID: PMC5607778.

79. Cambareri EB, Singer MJ, Selker EU. Recurrence of repeat-induced point mutation (RIP) in *Neurospora crassa*. Genetics. 1991;127(4):699–710. Epub 1991/04/01. PubMed PMID: 1827629; PubMed Central PMCID: PMCPMC1204397.

80. Fudal I, Ross S, Brun H, Besnard AL, Ermel M, Kuhn ML, et al. Repeat-induced point mutation (RIP) as an alternative mechanism of evolution toward virulence in Leptosphaeria maculans. Mol Plant-Microbe Interact. 2009;22(8):932–41. Epub 2009/07/11. doi: 10.1094/MPMI-22-8-0932. PubMed PMID: 19589069.

81. Wong S, Wolfe KH. Birth of a metabolic gene cluster in yeast by adaptive gene relocation. Nature Genetics. 2005;37(7):777–82. PubMed PMID: 15951822.

82. Orbach MJ, Farrall L, Sweigard JA, Chumley FG, Valent B. A telomeric avirulence gene determines efficacy for the rice blast resistance gene *Pi-ta*. Plant Cell. 2000;12(11):2019–32.

83. Rehmeyer C, Li W, Kusaba M, Kim YS, Brown D, Staben C, et al. Organization of chromosome ends in the rice blast fungus, *Magnaporthe oryzae*. Nucleic Acids Res. 2006;34(17):4685–701. PubMed PMID: 16963777.

84. Brown CA, Murray AW, Verstrepen KJ. Rapid expansion and functional divergence of subtelomeric gene families in yeasts. Curr Biol. 2010;20(10):895–903. Epub 2010/05/18. doi: 10.1016/j.cub.2010.04.027. PubMed PMID: 20471265; PubMed Central PMCID: PMC2877759.

85. Hirsch CD, Springer NM. Transposable element influences on gene expression in plants. Biochim Biophys Acta Gene Regul Mech. 2017;1860(1):157–65. Epub 2016/05/29. doi: 10.1016/j.bbagrm.2016.05.010. PubMed PMID: 27235540.

86. Chuong EB, Elde NC, Feschotte C. Regulatory activities of transposable elements: from conflicts to benefits. Nat Rev Genet. 2016;18(2):71–86. doi: 10.1038/nrg.2016.139. PubMed PMID: 27867194.

87. Lanver D, Müller AN, Happel P, Schweizer G, Haas FB, Franitza M, et al. The biotrophic development of *Ustilago maydis* studied by RNA-Seq analysis. Plant Cell. 2018;30(2):300–23. doi: 10.1105/tpc.17.00764. PubMed PMID: 29371439; PubMed Central PMCID: PMC5868686.

88. Kleemann J, Rincon-Rivera LJ, Takahara H, Neumann U, Ver Loren van Themaat E, van der Does HC, et al. Sequential delivery of host-induced virulence effectors by appressoria and intracellular hyphae of the phytopathogen *Colletotrichum higginsianum*. PLoS Pathog. 2012;8(4):e1002643. Epub 2012/04/13. doi: 10.1371/journal.ppat.1002643. PubMed PMID: 22496661; PubMed Central PMCID: PMCPMC3320591.

89. Soyer JL, El Ghalid M, Glaser N, Ollivier B, Linglin J, Grandaubert J, et al. Epigenetic control of effector gene expression in the plant pathogenic fungus *Leptosphaeria maculans*. PLoS Genet. 2013;10(3):e1004227. Epub 2014/03/08. doi: 10.1371/journal.pgen.1004227 PGENETICS-D-13-03255 [pii]. PubMed PMID: 24603691; PubMed Central PMCID: PMC3945186.

90. Jones P, Binns D, Chang HY, Fraser M, Li W, McAnulla C, et al. InterProScan 5: genome–scale protein function classification. Bioinformatics. 2014;30(9):1236–40. Epub 2014/01/24. doi: 10.1093/bioinformatics/btu031. PubMed PMID: 24451626; PubMed Central PMCID: PMC3998142.

91. Letunic I, Doerks T, Bork P. SMART: recent updates, new developments and status in 2015. Nucleic Acids Res. 2015;43(Database issue):D257–60. Epub 2014/10/11. doi: 10.1093/nar/gku949. PubMed PMID: 25300481; PubMed Central PMCID: PMC4384020.

92. Jorda J, Kajava AV. T-REKS: identification of Tandem REpeats in sequences with a K-meanS based algorithm. Bioinformatics. 2009;25(20):2632–8. Epub 2009/08/13. doi: 10.1093/bioinformatics/btp482. PubMed PMID: 19671691.

93. Zimmermann L, Stephens A, Nam SZ, Rau D, Kubler J, Lozajic M, et al. A completely reimplemented MPI bioinformatics toolkit with a new HHpred server at its core. J Mol Biol. 2018;430(15):2237–43. Epub 2017/12/21. doi: 10.1016/j.jmb.2017.12.007. PubMed PMID: 29258817.

94. Schardl CL, Young CA, Hesse U, Amyotte SG, Andreeva K, Calie PJ, et al. Plant-symbiotic fungi as chemical engineers: multi-genome analysis of the Clavicipitaceae reveals dynamics of alkaloid loci. PLoS Genetics. 2013;9(2):e1003323.

95. Kettles GJ, Bayon C, Canning G, Rudd JJ, Kanyuka K. Apoplastic recognition of multiple candidate effectors from the wheat pathogen *Zymoseptoria tritici* in the nonhost plant *Nicotiana benthamiana*. New Phytol. 2017;213(1):338–50. Epub 2016/10/04. doi: 10.1111/nph.14215. PubMed PMID: 27696417; PubMed Central PMCID: PMC5132004.

96. Bos JIB, Kanneganti T-D, Young C, Cakir C, Huitema E, Win J, et al. The C-terminal half of *Phytophthora infestans* RXLR effector AVR3a is sufficient to trigger R3a-mediated hypersensitivity and suppress INF1-induced cell death in *Nicotiana benthamiana*. Plant J. 2006;48(2):165–76. Epub 2006/09/13. doi: 10.1111/j.1365-313X.2006.02866.x. PubMed PMID: 16965554.

97. Ma LJ, van der Does HC, Borkovich KA, Coleman JJ, Daboussi MJ, Di Pietro A, et al. Comparative genomics reveals mobile pathogenicity chromosomes in *Fusarium*. Nature. 2010;464(7287):367–73. Epub 2010/03/20. doi: nature08850 [pii] 10.1038/nature08850. PubMed PMID: 20237561.

98. van Dam P, Fokkens L, Schmidt SM, Linmans JH, Kistler HC, Ma LJ, et al. Effector profiles distinguish formae speciales of *Fusarium oxysporum*. Environ Microbiol. 2016;18(11):4087–102. doi: 10.1111/1462-2920.13445. PubMed PMID: 27387256.

99. Inoue Y, Vy TTP, Yoshida K, Asano H, Mitsuoka C, Asuke S, et al. Evolution of the wheat blast fungus through functional losses in a host specificity determinant. Science. 2017;357(6346):80–3. Epub 2017/07/08. doi: 10.1126/science.aam9654. PubMed PMID: 28684523.

100. Yoshida K, Saitoh H, Fujisawa S, Kanzaki H, Matsumura H, Yoshida K, et al. Association genetics reveals three novel avirulence genes from the rice blast fungal pathogen *Magnaporthe oryzae*. Plant Cell. 2009;21(5):1573–91. Epub 2009/05/21. doi: 10.1105/tpc.109.066324. PubMed PMID: 19454732; PubMed Central PMCID: PMC2700537.

101. Ridout CJ, Skamnioti P, Porritt O, Sacristan S, Jones JD, Brown JK. Multiple avirulence paralogues in cereal powdery mildew fungi may contribute to parasite fitness and defeat of plant resistance. Plant Cell. 2006;18(9):2402–14. doi: 10.1105/tpc.106.043307. PubMed PMID: 16905653; PubMed Central PMCID: PMC1560912.

102. Houterman PM, Cornelissen BJ, Rep M. Suppression of plant resistance gene-based immunity by a fungal effector. PLoS Pathog. 2008;4(5):e1000061. Epub 2008/05/10. doi: 10.1371/journal.ppat.1000061. PubMed PMID: 18464895; PubMed Central PMCID: PMC2330162.

103. Selin C, de Kievit TR, Belmonte MF, Fernando WG. Elucidating the role of effectors in plant-fungal interactions: progress and challenges. Frontiers in microbiology. 2016;7:600. Epub 2016/05/21. doi: 10.3389/fmicb.2016.00600. PubMed PMID: 27199930; PubMed Central PMCID: PMC4846801.

104. Lu C, Chen J, Zhang Y, Hu Q, Su W, Kuang H. Miniature invertedrepeat transposable elements (MITEs) have been accumulated through amplification bursts and play important roles in gene expression and species diversity in *Oryza sativa*. Mol Biol Evol. 2012;29(3):1005–17. Epub 2011/11/19. doi: 10.1093/molbev/msr282. PubMed PMID: 22096216; PubMed Central PMCID: PMC3278479.

105. Fleetwood DJ, Scott B, Lane GA, Tanaka A, Johnson RD. A complex ergovaline gene cluster in epichloë endophytes of grasses. Appl Environ Microbiol. 2007;73(8):2571–9. PubMed PMID: 17308187.

106. van Dam P, Fokkens L, Ayukawa Y, van der Gragt M, Ter Horst A, Brankovics B, et al. A mobile pathogenicity chromosome in *Fusarium oxysporum* for infection of multiple cucurbit species. Sci Rep. 2017;7(1):9042. Epub 2017/08/24. doi: 10.1038/s41598-017-07995-y. PubMed PMID: 28831051; PubMed Central PMCID: PMC5567276.

107. Bolton MD, van Esse HP, Vossen JH, de Jonge R, Stergiopoulos I, Stulemeijer IJE, et al. The novel *Cladosporium fulvum* lysin motif effector Ecp6 is a virulence factor with orthologues in other fungal species. Mol Microbiol. 2008;69(1):119–36. Epub 2008/05/03. doi: 10.1111/j.1365-2958.2008.06270.x. PubMed PMID: 18452583.

108. Vargas WA, Sanz-Martín JM, Rech GE, Armijos-Jaramillo VD, Rivera LP, Echeverria MM, et al. A fungal effector with host nuclear localization and DNA-binding properties is required for maize anthracnose development. Mol Plant-Microbe Interact. 2016;29(2):83–95. Epub 2015/11/12. doi: 10.1094/MPMI-09-15-0209-R. PubMed PMID: 26554735.

109. Mesarich CH, Bowen JK, Hamiaux C, Templeton MD. Repeat-containing protein effectors of plant-associated organisms. Front Plant Sci. 2015;6:872. Epub 2015/11/12. doi: 10.3389/fpls.2015.00872. PubMed PMID: 26557126; PubMed Central PMCID: PMC4617103.

110. Grove TZ, Cortajarena AL, Regan L. Ligand binding by repeat proteins: natural and designed. Curr Opin Struct Biol. 2008;18(4):507–15. Epub 2008/07/08. doi: 10.1016/j.sbi.2008.05.008. PubMed PMID: 18602006; PubMed Central PMCID: PMCPMC3500881.

111. Johnson RD, Lane GA, Koulman A, Cao M, Fraser K, Fleetwood DJ, et al. A novel family of cyclic oligopeptides derived from ribosomal peptide synthesis of an in planta-induced gene, *gigA*, in *Epichloë* endophytes of grasses. Fungal Genet Biol. 2015;85:14–24. doi: 10.1016/j.fgb.2015.10.005. PubMed PMID: 26519220.

112. Wosten HA, Bohlmann R, Eckerskorn C, Lottspeich F, Bolker M, Kahmann R. A novel class of small amphipathic peptides affect aerial hyphal growth and surface hydrophobicity in *Ustilago maydis*. EMBO J. 1996;15(16):4274–81. Epub 1996/08/15. PubMed PMID: 8861956; PubMed Central PMCID: PMC452153.

113. Teertstra WR, van der Velden GJ, de Jong JF, Kruijtzer JA, Liskamp RM, Kroon-Batenburg LM, et al. The filament-specific Rep1-1 repellent of the phytopathogen *Ustilago maydis* forms functional surface-active amyloid-like fibrils. J Biol Chem. 2009;284(14):9153–9. Epub 2009/01/24. doi: 10.1074/jbc.M900095200. PubMed PMID: 19164282; PubMed Central PMCID: PMCPMC2666566.

114. Christensen MJ, Bennett RJ, Ansari HA, Koga H, Johnson RD, Bryan GT, et al. *Epichloë* endophytes grow by intercalary hyphal extension in elongating grass leaves. Fungal Genet Biol. 2008;45:84–93.

115. Voisey CR. Intercalary growth in hyphae of filamentous fungi. Fungal Biol Rev. 2010;24:123–31.

116. Bultman TL, Leuchtmann A. Biology of the Epichloë-*Botanophila* interaction: an intriguing association between fungi and insects. Fungal Biol Rev. 2009;22:131–8.

117. Stewart EL, McDonald BA. Measuring quantitative virulence in the wheat pathogen *Zymoseptoria tritici* using high-throughput automated image analysis. Phytopathology. 2014;104(9):985–92. Epub 2014/03/15. doi: 10.1094/PHYTO-11-13-0328-R. PubMed PMID: 24624955.

118. Stewart EL, Hagerty CH, Mikaberidze A, Mundt CC, Zhong Z, McDonald BA. An improved method for measuring quantitative resistance to the wheat pathogen *Zymoseptoria tritici* using high-throughput automated image analysis. Phytopathology. 2016;106(7):782–8. Epub 2016/04/07. doi: 10.1094/PHYTO-01-16-0018-R. PubMed PMID: 27050574.

119. Wessling R, Epple P, Altmann S, He Y, Yang L, Henz SR, et al. Convergent targeting of a common host protein-network by pathogen effectors from three kingdoms of life. Cell host & microbe. 2014;16(3):364–75. Epub 2014/09/12. doi: 10.1016/j.chom.2014.08.004. PubMed PMID: 25211078; PubMed Central PMCID: PMC4191710.

120. Win J, Chaparro-Garcia A, Belhaj K, Saunders DG, Yoshida K, Dong S, et al. Effector biology of plant-associated organisms: concepts and perspectives. Cold Spring Harbor symposia on quantitative biology. 2012;77:235–47. Epub 2012/12/12. doi: 10.1101/sqb.2012.77.015933. PubMed PMID: 23223409.

121. Heath MC. Nonhost resistance and nonspecific plant defenses. Curr Opin Plant Biol. 2000;3(4):315–9. PubMed PMID: 10873843.

122. Derevnina L, Dagdas YF, De la Concepcion JC, Bialas A, Kellner R, Petre B, et al. Nine things to know about elicitins. New Phytol. 2016;212(4):888–95. Epub 2016/09/02. doi: 10.1111/nph.14137. PubMed PMID: 27582271.

123. Du J, Verzaux E, Chaparro-Garcia A, Bijsterbosch G, Keizer LC, Zhou J, et al. Elicitin recognition confers enhanced resistance to *Phytophthora infestans* in potato. Nature plants. 2015;1(4):15034. Epub 2015/01/01. doi: 10.1038/nplants.2015.34. PubMed PMID: 27247034.

124. Dupont PY, Eaton CJ, Wargent JJ, Fechtner S, Solomon P, Schmid J, et al. Fungal endophyte infection of ryegrass reprograms host metabolism and alters development. New Phytol. 2015;208(4):1227–40. doi: 10.1111/nph.13614. PubMed PMID: 26305687.

125. Becker M, Becker Y, Green K, Scott B. The endophytic symbiont *Epichloë festucae establishes an epiphyllous net on the surface of Lolium perenne* leaves by development of an expressorium, an appressorium-like leaf exit structure. New Phytol. 2016;211:240–54. doi: 10.1111/nph.13931. PubMed PMID: 26991322.

